# Resolving complex structures at oncovirus integration loci with conjugate graph

**DOI:** 10.1101/2020.05.17.100578

**Authors:** Wenlong Jia, Chang Xu, Shuai Cheng Li

**Author notes:** These authors contributed equally to this work (Wenlong Jia, Chang Xu).

## Abstract

Oncovirus integrations cause copy number variations (CNVs) and complex structural variations (SVs) on host genomes. However, the understanding of how inserted viral DNA impacts the local genome remains limited. The linear structure of the oncovirus integrated local genomic map (LGM) will lay the foundations to understand how oncovirus integrations emerge and compromise the host genome’ s functioning. We propose a conjugate graph model to reconstruct the rearranged local genomic map at integrated loci. Simulation tests prove the reliability and credibility of the algorithm. Applications of the algorithm to whole-genome sequencing data of Human papillomavirus (HPV) and hepatitis B virus (HBV)-infected cancer samples gained biological insights on oncovirus integrations. We observed five affection patterns of oncovirus integrations from the HPV and HBV-integrated cancer samples, including the exon loss, promoter gain, hyper-amplification of tumor gene, the viral cis-regulation inserted at the single intron and at the intergenic region. We found that the focal duplicates and host SVs are frequent in the HPV-integrated LGMs, while the focal deletions and complex virus SVs are prevalent in HBV-integrated LGMs. Furthermore, with the results yields from our method, we found the enhanced microhomology-mediated end joining (MMEJ) might lead to both HPV and HBV integrations, and conjectured that the HPV integrations might mainly occur during the DNA replication process. The conjugate graph algorithm code and LGM construction pipeline, available at https://github.com/deepomicslab/FuseSV.

**Key points:**

- Conjugate graph model resolves complex local genomic strcuture at oncovirus integration loci.
- Local genome maps reveal five affection patterns of oncovirus integrations.
- Microhomology bases and small insertions are enriched at the junctions of structural variations and virus integrations.
- HPV and HBV integrations may be induced by the enhanced microhomology-meditated mechanism.

## Introduction

Oncovirus integrations (VITs) could induce cancer genome instability [1–5], emerging as copy number variations (CNVs) and complex structural variations (SVs). Adjacent genes to the integrated sites are frequently dysregulated, such as *TERT* in hepatocellular carcinoma [5, 6] and *MYC* in cervical carcinoma [7, 8]. However, the explicit structure of the rearranged local genomic map (LGM) remains elusive. Some studies have reported the local maps at the integration site of human papillomavirus (HPV). Adey et al. resolved the haplotype of the HeLa cell line and HPV18-integrated genomic locus by consolidating results from a range of sequencing technologies, including the whole-genome shotgun sequencing, the fosmid mate-pair library with large insert size, and PacBio single-molecule sequencing data [1]. Akagi *et al*. constructed manually preliminary LGMs flanking integration sites in HPV-positive samples [2], including head and neck squamous cell carcinomas (HNSCC) and cervical cancers. However, oncovirus integrations demand the resolved local maps systematically, which would lead to more insights. Accurate LGMs analysis lays the foundations to understand how oncovirus integrations emerge and compromise the host genome’s functioning. A computational model and algorithm are necessary to obtain the local genomic maps.

Although a growing number of studies examined the oncovirus integration effects on the expression regulations and genome stability in the flanking host genomic regions [6–13], our understanding of how inserted viral DNA influences the local genome remains limited. The LGMs can provide the haplotyping information to promote the investigation, such as the pairwise integrations, host and viral rearrangements, inserted viral regulatory elements, and the copy number changes. Further, the LGM linear structure can facilitate the viral integration mechanism research with junction breakpoints phased together, as well as the alternative splicing and virus-host fusions. Several factors hinder the LGMs accurate construction, including the sequencing fluctuations, short genomic segments, repeated elements, and insufficient junction reads. Here, we introduce a graph-based algorithm to overcome these difficulties and construct LGMs at the oncovirus integration sites from cancer whole genome sequencing (WGS) data. Using multiple real and *in silico* datasets, we show that the algorithm is able to construct credible LGMs of oncovirus integrations and complex rearrangements on cancer genome. The LGM results further facilitate our investigation on the regulatory effects and formation mechanism accompanied by virus integrations.

## Methods

### Conjugate graph

We first consider the problem of reconstructing the disrupted genome with perfect NGS data and breakpoint detection, giving rise to no ambiguity. Based on the breakpoints, we derive two sets of reference genomic regions respectively from within (1) the human/host genome (denoted H = {*h*_1_, *h*_2_, … , *h*_m_}), and (2) the virus genome (denoted *V* = {*v*_1_, *v*_2_, … , *v*_m_}). Each region *s* ∈ *H* ∪ *V* can be either on the positive or the negative DNA chain, denoted *s*^+^ and *s*^−^ respectively. Associated with each region *s*is a number *c*(*s*) which indicates the copy number of the region.

We obtain from the NGS data a set *J* of junctions, each of which connects two regions in *H*∪ *V*. The junctions includes endogenous SVs on human and virus genome, VITs, and reference segment connections. Each μ ∈ J is an ordered pair 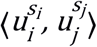, where *s*_*i*_, *s*_*j*_ ∈ {+, −} and *u*_*i*_,*u*_*j*_ ∈ *H* ∪ *V*. As DNA possesses a double helix structure, a junction segment is equivalent to its reverse complementary segment. Therefore, two junction segments 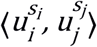 and 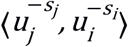 are equivalent, where −*s*_i_ denotes the reverse complementary chain of *s*_i_. Note that junction segments are directed, that is, 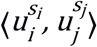 is different from 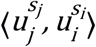. Meanwhile, each μ ∈ J may also occur multiple times in NGS data, denoted by *c*(μ), indicating the copy number of sequencing reads (i.e., split-reads) crossing a junction.

A directed graph *G*= (*U*, *E*) is hence constructed from the NGS data, where the vertices *U*= {*u*^+^, *u*^−^ | *u*∈ *H* ∪ *V*} and edges *E*= J. Given that H, V, J, and c contain no error, then

- G is connected.
- 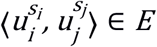 if and only if 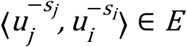, and
- a path that visits each vertex *u* ∈ *U* exactly *c*(*u*) times, and each edge exactly *c*(μ) times exists.

We refer to 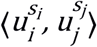 and 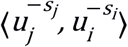 as conjugate edges, and the vertices *u*^+^ and *u*^−^ as conjugate vertices. We call the resultant graph a conjugate graph.

### Local genomic map

Local genomic map (LGM) is defined as a path over the conjugate graph, where each *u* ∈ *H* ∪ *V* is visited exactly *c*(*u*) times, and each μ ∈ J is traversed exactly *c*(μ) times. Hence an LGM of length *l* gives us a sequence of visited vertices 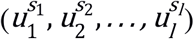 where *u*_*i*_ ∈ *H* ∪ *V* and *s*_*i*_ ∈ {+, −}. Without loss of generality, we assume that both the starting and ending point of an LGM are from the positive chain of the host, that is, 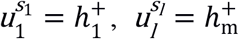. Note that we assume the path is along the positive strand. If the local genomic map contains a vertex of a negative strand, it implies an inversion event.

### Existence of LGM in the conjugate graph

We show that it is possible to know if a conjugate graph G contains an LGM. Define the in-copy of a vertex *u*^*s*^ as the total copy number of edges with *u*^*s*^ as *target* vertex; that is, 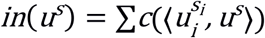. Similarly, we can define 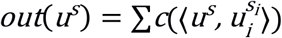.

Clearly, if G contains an LGM, then it satisfies the following properties:

1. *c*(*u*) = *in*(*u*^−^) + *out* (*u*^+^) = *out*(*u*^−^) + *in*(*u*^+^), ∀ *u* ∈ *H* ∪ *V* – {*h*_1_, *h*_m_}
2. 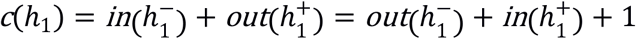 and 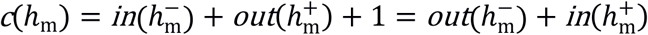
3. A path exists from 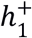 to *u*^+^ or *u*^−^, and a path exists from *u*^+^ or *u*^−^ to 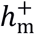.

Note that, to form a path, we add a putative edge links *sink* vertex back to *source* vertex (property (2)). We refer to the properties (1) and (2) as degree balance, and property (3) as reachability.

An LGM of a conjugate graph shares similarities with an Eulerian path of an Eulerian Graph. The differences lie in that the vertices and edges are conjugated in LGMs. We adopt an algorithm derived from the Hierholzer’s algorithm to find LGMs.

Similar to Hierholzer’s algorithm, the algorithm for finding LGMs starts from an arbitrary vertex 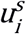, and recursively follow a trail of out-going edges from the vertex until returning to the vertex 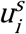 itself, forming a circuit *C*. Due to the conjugation property, after we traverse an edge 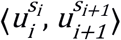, the next edge with unused copy numbers we can choose from is either in the form of 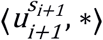 or 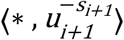, where * represents a wildcard. Similar to the Eulerian path argument, as long as we have a vertex in the graph with unused copy numbers, the degree balance property ensures that it is always possible to find a circle that starts from and ends in the vertex. Therefore, given a vertex 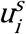 in the circle C that still has unused copy numbers, we can find a new circle C’ which can be merged with C to form a longer circular path. The above process is performed iteratively until we drain the copy numbers in the conjugate graph.

#### Theorem 1

A conjugate graph contains a local genomic map if and only if vertex 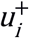 or 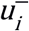 of any region *u* ∈ *H* ∪ *V* is in a path from 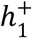 to 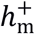 or a path from 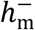 to 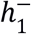, and furthermore, *u* satisfies the degree balance condition.

### Correcting the copy numbers

In the case of imperfect NGS data, the estimated copy numbers may contain fluctuations. The conjugate graph constructed would then lead to erroneous LGMs or no LGMs at all. To prevent such cases, we perform corrections to the graph under the principle that a corrected graph would contain an LGM, that is, the resultant conjugate graph should fulfill the two properties of reachability and degree balance.

It is possible to have missing edges or vertices due to reasons such as low sequencing depth and failure in detecting breakpoints which could break the property of reachability. To solve this in a parsimonious manner, we only consider the insertion of additional edges that are likely to exist in a normal genome, such as 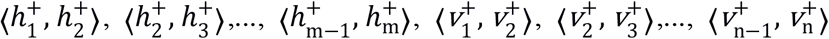, and 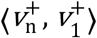 if the virus in question has a circular DNA structure.

We propose an integer linear programming approach to address the degree balance property. Assign each region *u* and each junction μ to a target copy number *t*(*u*) or *t* (μ) (Equations 1d and 1e), respectively, to satisfy the degree balance property (Equation 1b). The objective is to minimize the disagreement between the observed copy number and the target copy number (Equation 1a).

minimize

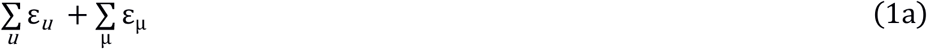

subject to

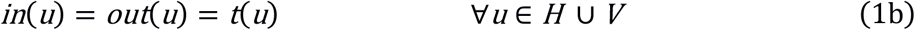

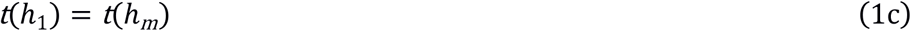

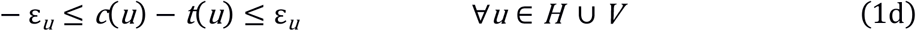

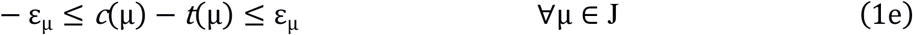

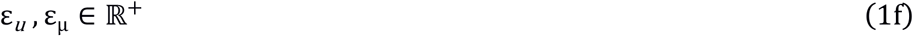

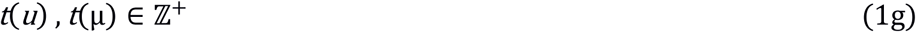

Additional domain knowledge can be incorporated to further refine the linear programming process. For instance, the copy number of a virus DNA segment can be easily overestimated due to the existence of free virus in cells and other viral integration sites, and the copy number of DNA segments too short can be underestimated due to alignment difficulties. In those circumstances, we may assign lower penalty to the change of their copy numbers during the integer linear programming. The virus can have much more copy number than tumor genomes, due to the free virus genome and other viral integration sites. Considering this point, our model does not ask the copy numbers of virus genomic segments to be fully used in the LGMs. In our model, we assigned small weights to the viral segments, which allows excessive virus copies out of the current LGM if necessary.

### Finding LGM

Our basic algorithm identifies an LGM path by iteratively selecting edges with non-zero weight at random to find circuits (called unit-cycles) and merge them. When determining the order of unit-cycles to be merged, the basic algorithm has no preference for any one possible LGM over the others. Principles of parsimony can be adopted; for instance, reference edges are preferred over edges that cross SVs or VITs breakpoints.

### Simulation evaluation

One subtype (B2) of HBV and two subtypes (HPV16 and HPV18) of HPV are chosen to simulate virus integrations. Virus reference genome was randomly divided into segments. Integration sites in human reference genome (GRCh37) were randomly chosen inside reported integration hotspot regions (near gene TERT, CCNE1, KMT2B for HBV, LRP1B, FHIT for HPV16 and at the upstream of MYC for HPV18). Lengths of segments are random values within empirical ranges based on our research (2000-35000bp for each human segment, 1000-4000bp and 500-2000bp for each HPV and HBV segment respectively). Human and virus segments were then concatenated with regard to the different integration modes (**Supplementary Figure S1**) and saved as FASTA format files.

Simulated sequencing was performed using SAMtools (v1.3) wgsim utility at 30X average depth and 150bp read length [14]. Lower average depth (15X) is also introduced to complex integration cases to test algorithm’s sensitivity. Since wgsim simulates all bases with the same quality score, we have also replaced the quality value of each sequence with qualities from actual 150bp sequencing data to resemble real sequencing results. Simulated sequencing reads are aligned back to human reference genome. Depth of each segment is calculated using SAMTools (v1.3, depth command) on uniquely mappable regions. Sequence mappability is determined by performing a 150bp read length simulated sequencing and aligning back to human genome. Regions covered by multiple-mapped reads are excluded in depth calculation.

All simulated samples are considered to have homogeneous integrations. Integration breakpoints are called by FuseSV. Structural variations on human and virus genome are called by Meerkat [15] and seeksv [16]. Local genomic map of all simulated cases are recovered by the algorithm. Complex cases can have multiple possible LGMs.

### Evaluation of integer linear programming

Test were performed to evaluate the robustness of the linear programming approach. We chose six simulated samples with large segment counts and tandem duplications, and then randomly assign fluctuations to segments’ depth. The fluctuation values are within [− *c_avg_*/2, *c_avg_*/2] where *c*_*avg*_ is the average haploid sequencing depth. The randomization is done 1 times for each chosen samples and we performed linear programming for each iteration. 98% of all 6 cases are corrected to our expected copy number profile. The other 2% cases had unit-cycles with one more or one less copy, still representing the correct LGM structure.

### WGS data analysis

Tumor samples from cell lines have no corresponding normal sample. From our data repository, we selected a normal sample with no CNV at known integration regions. WGS data was aligned to human reference genome (GRCh37) using BWA [17]. HPV and HBV integrations are detected by FuseSV. Meerkat [15] and seeksv [16] are used to call structural variations on human and viral genome. CNV profile, tumor purity and ploidy are generated by Patchwork [18]. Segment depths were normalized against GC bias by deepTools [19].

### Copy number calculation of segments and junctions in LGM

In tumor and control samples, the average depth of each host segment was calculated from GC-bias corrected alignments using SAMtools depth utility with minimum base and mapping quality set as 5 and 1, respectively. Call the purity of tumor-cell in tumor sample as *Purity*_*t*_, the average ploidy of pure tumor-cell in tumor sample as *Ploidy*_*t*_, both *Purity*_*t*_ and *Ploidy*_*t*_ are from Patchwork result [18]. And the average ploidy of pure normal cell as *Ploidy*_*n*_ (assumed as 2). The average ratio of the DNA from normal cells (*Ratio*_*n*_) in each genomic region was calculated as below:

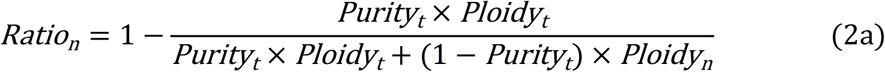

For each host segment, the germline copy ratio of control normal sample (*CopyRatio*_*n*_) can be obtained from WGS data (generally is one). The depth of this segment from pure tumor-cell in tumor sample (*Depth*_*t*_) could be obtained from the observed segment average depth (*Depth*_*o*_) and the whole genome average depth (*Depth*_*g*_) via formula

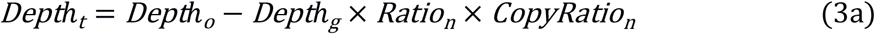

To calculate the copy number of each host segment, the haploidy depth (*HaploDepth*_*t*_) corresponding to single copy (CN=1) in the tumor cell could be obtained via formula

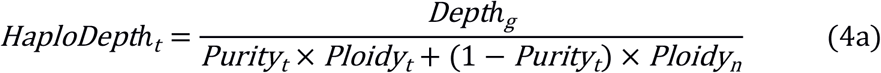

The copy number of each host segment is obtained from its depth (*Depth*_*t*_) divided by the *HaploDepth*_*t*_. Similarly, the copy numbers of VITs and SVs are calculated from the split-reads count divided by the *HaploDepth*_*t*_. The copy numbers of viral segments are also obtained from the viral *Depth*_*t*_ divided by the *HaploDepth*_*t*_. Note that the *Depth*_*t*_ of viral segment is calculated via formulas (2a) and (3a) with both *CopyRatio*_*n*_ and *Ploidy*_*n*_ set as zero, as we assumed all viral segments are reserved in tumor cells. The major and minor copy numbers (integer) of LGM region are determined from Patchwork CNV result [18], where we can get the objective copy number (integer) of the bilateral segments (source and sink) of LGM.

### Construction of virus genome in each sample

We extract the unmapped and soft-clipped reads from alignment results of WGS and realign these reads to HPV and HBV references from the NCBI nucleotide database. The viral subtype with the most uniquely aligned reads and coverage of at least 20% was selected as the major one. Mutations (SNV and InDel) are detected in an iterative process to amend the viral genome sequence until no more mutations can be identified. This procedure is also adopted in our previous study [4]. The constructed viral genome is applied in the subsequent viral integration and LGM analyses of the corresponding cancer sample. Two reads types (span-reads and split-reads) are sought to support the viral integrations.

### Microhomology at the junction sites

Microhomology bases (MHs) are detected at the junction sites of viral integrations and structural variations. Two types of MH bases are considered: the aligned and slipped MHs. Basically, the aligned microhomology is vertically matched bases covering or flanking the junction site, and the slipped microhomology has offset between the matched bases [20]. The slipped MH bases are required to have ≥ 3*bp* size and at least one base overlap. Criteria to determine the MH existence: i) the MH bases (≥ 2*bp*) must cover or locate next to the junction site, or ii) locate in flanking region (≤ 5*bp*) of the junction site and the MH size must be ≥ 4*bp* according our previous finding [7], or iii) merge 1bp-gaped neighbor MH bases and subject to the second criterion.

## Results

### Conjugate graph to construct local genomic map

Based on the breakpoints of VITs and SVs, both the host local genome and viral genome are divided into segments (**Figure 1A-B**), as vertices in the graph model (**Figure 1D**). Copy numbers (CNs) of segments are as the weights of vertices. Junctions of SVs and VITs are modeled as directed edges connecting two vertices, and their split-reads counts are converted to CN as the weights of the edges (**Figure 1C-D**). Integer linear programming is applied to correct weights of vertices and edges to reduce segment depth fluctuations, as well as compliment insufficient split-reads. As genome DNA has double strands, the vertices or edges are conjugated, i.e, a vertex or an edge and their reverse complementary counterparts and share the respective weights. We then use an algorithm modified from the Eulerian Circuits traversal rule to find available circular paths (called unit-cycles, **Figure 1E**), and then merge them to construct the local genomic map consisting of host and virus segments (**Figure 1F**).

**Figure 1.**
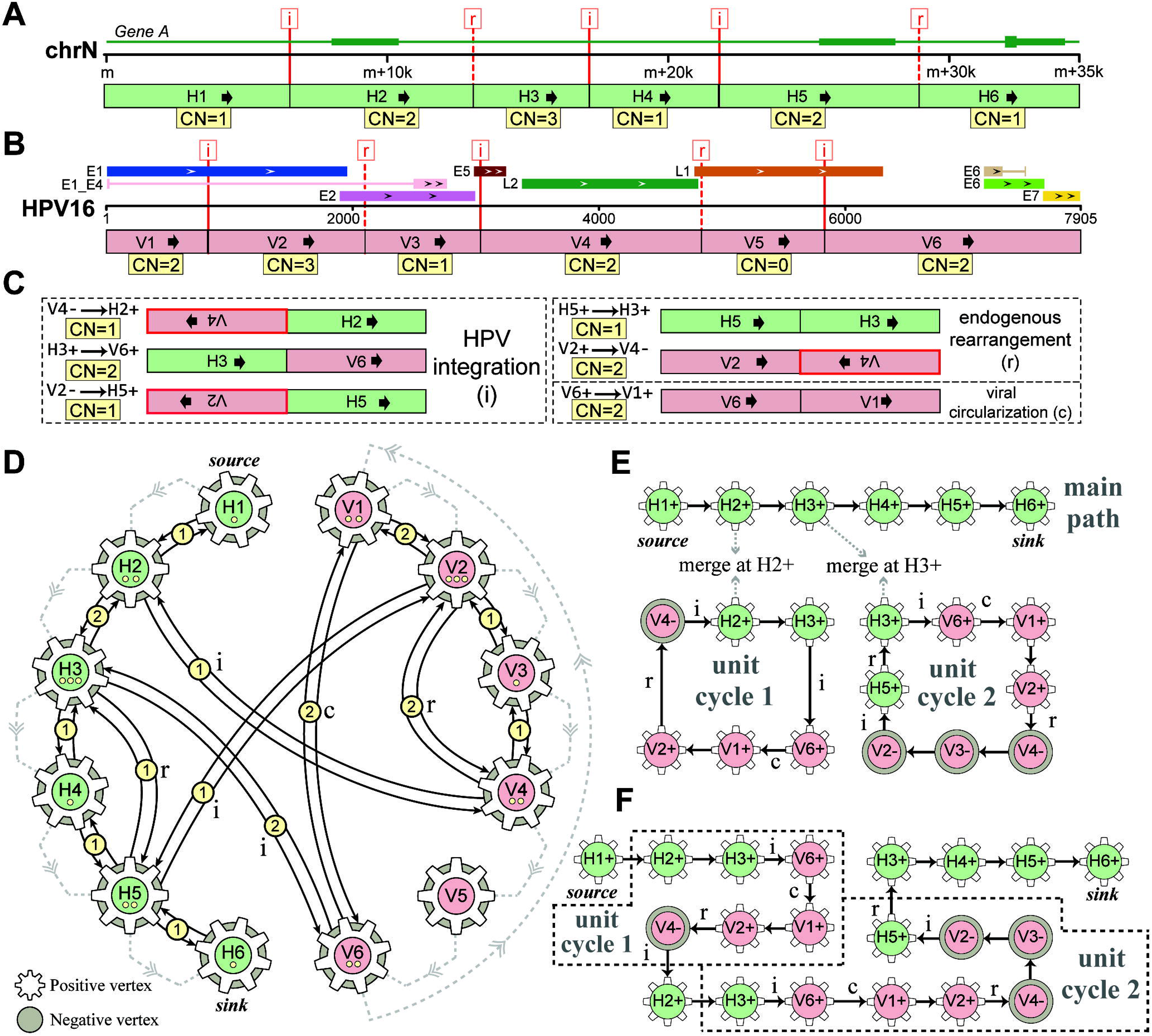
Demo representation of Conjugate graph resolved LGM of HPV16 integrations. Breakpoints of SVs (‘r’) and VITs (‘i’) are represented by dashed and solid red lines respectively, by which the host local (**A**) and HPV16 (**B**) genomes are divided into segments (host: H1~H6; HPV: V1~V6) denoted with copy number (CN, yellow filled frame) adjusted by integer linear programming (ILP). (**C**) Variant segment junctions utilized in Conjugate graph, including VITs (‘i’), SVs (‘r’) and viral circularization (‘c’). Junctions are denoted as pairwise segment ID and strand with copy number adjusted by ILP. Note that segments in red frame are reverse complementary counterpart. (**D**) The Conjugate graph model with directed vertices and edges representing segments and junctions respectively. The conjugate property of vertex (positive and negative) and junction (bi-direction) corresponds to the double strands of DNA. Dots on vertices and circled numbers on edges denote the copy number adjusted by ILP. Dashed edges indicate the segment junctions along the reference genome. (**E**) The unit-cycle paths resolved from the Conjugate graph. The main path extends from the source vertex along non-zero weighted edges to the sink vertex. (**F**) The LGM after merging unit-cycles with the main path.

In our algorithm, the first upstream and last downstream host segments are called the source and sink vertices of the conjugate graph (**Figure 1D-E**) respectively. The algorithm first finds the main path, which starts from the source vertex and extends along the edges with non-zero weight until it reaches the sink vertex (**Figure 1E**). If a vertex or an edge is included in the path, its weight on the graph decreases by one.

Then, starting at a vertex with non-zero weight left, our algorithm forms a unit-cycle by randomly walking along edges with non-zero weight until the vertex itself has reached again. More unit-cycles can be found iteratively until the all the weights are drained. Finally, all unit-cycles are merged with the main path to get putative LGMs. Due to the random walking and merging of the unit-cycles, there could be multiple LGM candidates generated from the conjugate graph, and our algorithm will report all possible LGMs.

### Evaluation on in silico data

To evaluate the reliability of the conjugate graph algorithm, we simulated a set of oncovirus integration LGMs at genomic loci of well-known hotspot genes, including HBV integrations (TERT, CCNE1, and KMT2B) [11] and HPV integrations (LRP1B, FHIT, and MYC) [7]. Combining the different SV types, copy numbers, and viral segments, we generated eight LGM simulated modes in each gene locus, such as substitutive integrations, inversed insertions, and complex tandem duplications (**Supplementary Figure S1**). In total, we prepared simulated sequencing data of 48 LGM cases for evaluation (**Supplementary Table S1**). The conjugate graph algorithm successfully recovered all simulated LGMs at HBV and HPV integrated sites. The integer linear programming obtained the copy numbers of all segments and junctions (SVs and VITs) with zero offset from the expected values. Moreover, for complex structures with high CN segments, the algorithm reported other possible candidate genomic maps that also satisfy the observed conjugate graph. The performance on in silico data supports the reliability of the conjugate graph to resolve diverse local genomic maps.

### On WGS data of HPV-integrated cervical cancers

We first applied the developed method to reconstruct the LGMs at HPV integration loci in four cervical carcinoma samples (T4931, T6050, HeLa, and SiHa) [7]. All samples have arm-level copy number changes of the HPV-integrated chromosomes (**Supplementary Figure S2**). The conjugate graph algorithm constructed LGMs of all validated HPV integrations with most CN offsets less than one (73.3%, **Supplementary Table S2** and **S3**).

First, the LGM on unbalanced polyploidy was successfully constructed in sample T4931. The HPV16 integrations locate at gene GLI2 locus on chr2q (heterozygous pentaploid, minor CN = 2, **Supplementary Figure S2A**). As the integration might locate on the paternal or maternal chromosomes, our method algorithm provides two LGM candidates (**Figure 2** and **Supplementary Figure S3**). The resolved LGM consists of two unit-cycles containing three virus integrations (VITs), virus circularization (V-CIRC), one viral inversion, and one host tandem duplication (**Figure 2C**). The HPV segments carry complete viral long control region (LCR) and insert at the intron region of gene GLI2.

**Figure 2.**
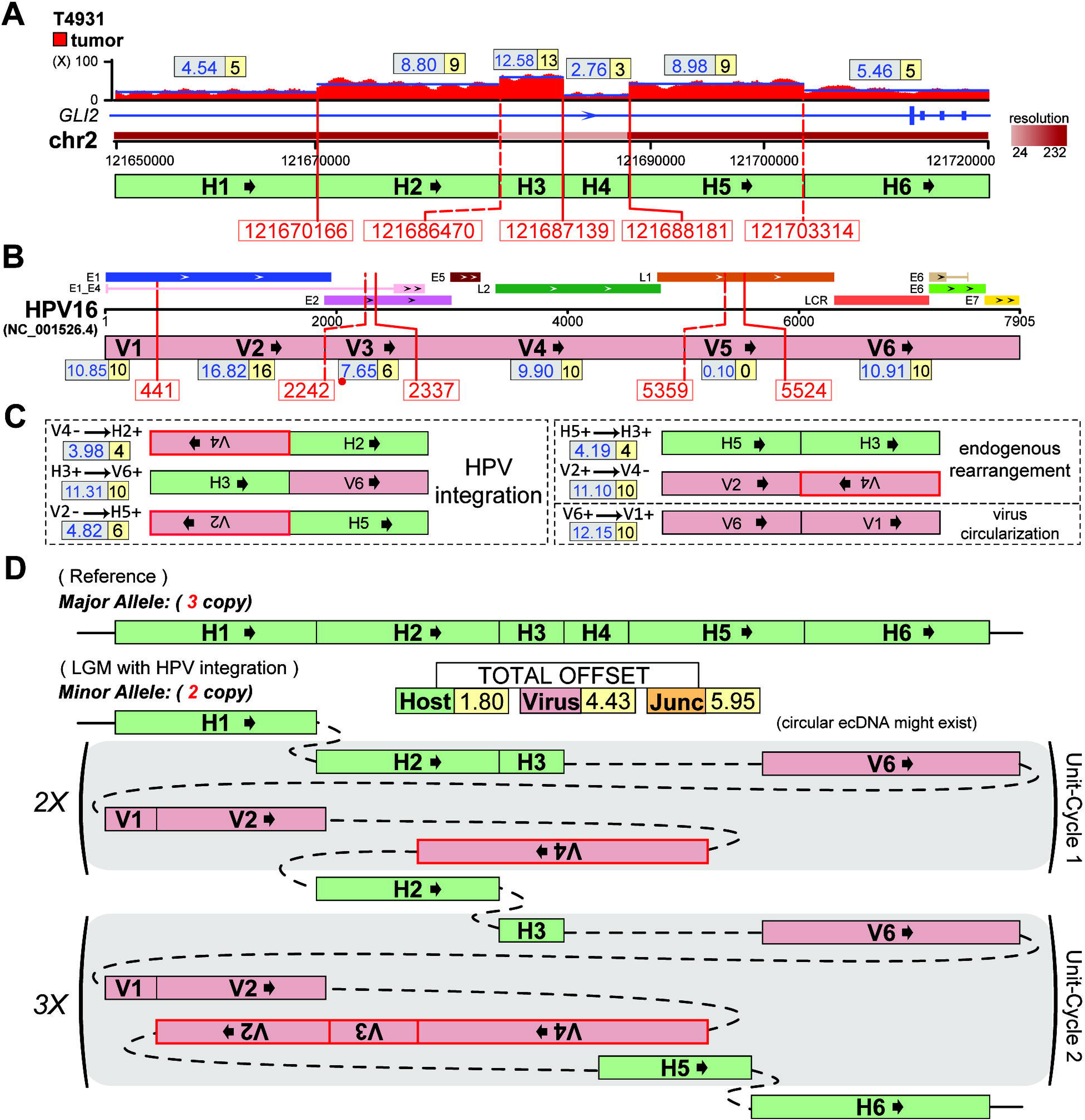
Presentative LGM at HPV integration sites (gene GLI2) on chr 2 (minor allele) of the T4931 sample [7]. (**A**) Human genomic region flanking HPV integrations are divided into six segments (H1~H6, in different resolutions) by VITs (red solid-line) and SVs (red dashed-line) denoted with breakpoints. Depth spectrum is displayed with original (grey frame) and ILP-adjusted (yellow frame) copy numbers of segments. (**B**) Segmentation (V1~V6) of HPV16 genome by VITs and SVs. The segment (V3) less than 100bp is marked with red dot. (**C**) Variant segment junctions utilized in Conjugate graph. (**D**) Resolved alleles of the ‘Simplest LGM’ are indicated as string of coloured segments with copy times, including reference allele (major allele) and that harbours HPV16 integrations (minor allele). Sum of absolute offset of segments and junctions are shown. Unit-cycles (shadowed areas) in LGM are denoted with repeat time. Note that the unit-cycles might exist as circular extrachromosomal DNA (ecDNA).

Second, we solved the LGM with asymmetric copy of the source and sink vertices. The chr8q of HeLa cell line shows different copy numbers at the HPV18 integrated sites: the upstream and downstream are tetraploid and hexaploid respectively, both of which are heterozygous (minor CN = 1, **Supplementary Figure S2B**). Previous study has reported that the HPV18 integrates on the major allele of the tetraploid [1]. The constructed LGM contains three unit-cycles consisting of four VITs and the V-CIRC (**Supplementary Figure S4**). The unit-cycles are evenly distributed in the three copies of the major allele. The LGM structure is similar to the reported [1], but with different copy numbers and breakpoints of the unit-cycles.

Third, two LGMs appeared as diploid with loss of heterozygosity (LOH). Both of the T6050 sample and SiHa cell line have HPV16 integrations at chr13q (LOH, **Supplementary Figure S2C-D**). The LGM of T6050 contains one unit-cycle consisting of two VITs, the V-CIRC, and one host tandem duplication at the gene KLF12 locus (**Supplementary Figure S5**). Interestingly, this unit-cycle has different copy times (two and three) in the remained diploid alleles, implying the possible existence of circular extrachromosomal DNA (ecDNA). The SiHa cell line has two unit-cycles in the LGM at the HPV16 integration sites (downstream of the gene KLF12, **Supplementary Figure S6**). The unit-cycles contain two VITs, the V-CIRC, one viral deletion, and one host deletion. The structure of the unit-cycles is the same as in the previous reports [2].

### On WGS data of HPV-integrated HNSCC cancers

We then applied the conjugate graph algorithm on two HPV16-positive HNSCC cell lines (UM-SCC-47 and UPCI-SCC090) [2]. The resolved HPV16-integrated LGMs are more complex than the four cervical cancers. Different from the published preliminary results [2], our method reported all detected genomic segments and breakpoints.

The first LGM contains tandem duplication of partial gene body of tumor suppressor, and might lead to the truncation of the coding-frame. The UM-SCC-47 cell line has HPV16 integrations at the tumor suppressor TP63 locus on chr3q [21, 22]. The broad region (3q26.31-q29) shows heterozygous triploid (minor CN = 1), and the focal region (in q28) at the HPV16 integrations has prominent high copy number gains (**Supplementary Figure S7A**). In total, six HPV16 integrations and four endogenous rearrangements are considered to construct two LGMs (**Supplementary Table S4**) corresponding to the minor (**Supplementary Figure S8**) and major alleles (**Supplementary Figure S9**). Our algorithm successfully obtained five unit-cycles. Interestingly, all segments of the NO.1 unit-cycle are from the host genome and form a tandem duplication structural variation with an inner deletion (the ‘H3’ segment). The NO.3 and NO.4 unit-cycles both have high copy numbers (5 and 49 respectively) and mainly contribute to the focal gain on the host genome. The minor allele and major allele LGMs share the same unit-cycles and differ only in copy numbers.

Another LGM has focal hyper-amplification inducing the copy number gain of oncogene. The UPCI-SCC090 cell line has HPV16 integrations in chr6p21.2. The chr6 is heterozygous triploid (minor CN = 1), and the HPV-integrated focal region shows a higher copy number (**Supplementary Figure S7B**). The focal host region contains seven HPV16 integrations and six endogenous rearrangements. These breakpoints divide the host and virus genome into a total of 28 segments (**Supplementary Table S4**), significantly more than the other samples mentioned above. Two LGMs are constructed for virus integrations on the minor (**Supplementary Figure S10**) and major allele (**Supplementary Figure S11**), respectively. The minor allele LGM is formed by seven unit-cycles. The NO.1 unit-cycle crosses the host and viral genomes five times with inversions and integrations. The NO.6 and NO.7 unit-cycles mainly contribute to the high copy number of this focal region with 19 and 14 copies, respectively. The major allele LGM has a similar global structure but one more unit-cycle. This LGM involves multiple genes, such as PI16, FGD2, PIM1, and SNORD112. The proto-oncogene PIM1 is completely located in the ‘H9’ segment with a high copy number (CN = 58). In addition, after the LGM construction, the remaining HPV16 genome copies formed two types of free viral genome by our conjugate graph algorithm: the complete (CN = 655) and with deletion (CN = 22). Moreover, the remaining host segments might exist as the ecDNAs.

### On WGS data of HBV-integrated hepatocellular carcinomas

Next, we applied the conjugate graph algorithm on the HBV-infected hepatocellular carcinomas (HCC). Six HCC samples were selected from the previous study that reported the genome-wide detection of HBV integrations [11]. Four samples have arm-level copy number changes of the HBV-integrated chromosomes, but none shows the focal gain at the integration sites (**Supplementary Figure S12**). The constructed LGMs further reveal that most of the inserted HBV DNA segments replace the host segments and consequently lead to the deletions on the human genome (**Supplementary Table S5** and **S6, Figure 3, Supplementary Figure S13-S17**), different from the tandem duplications induced by the HPV integrations. Additionally, the remaining HBV segment copies imply the free viral genomes in five samples.

**Figure 3.**
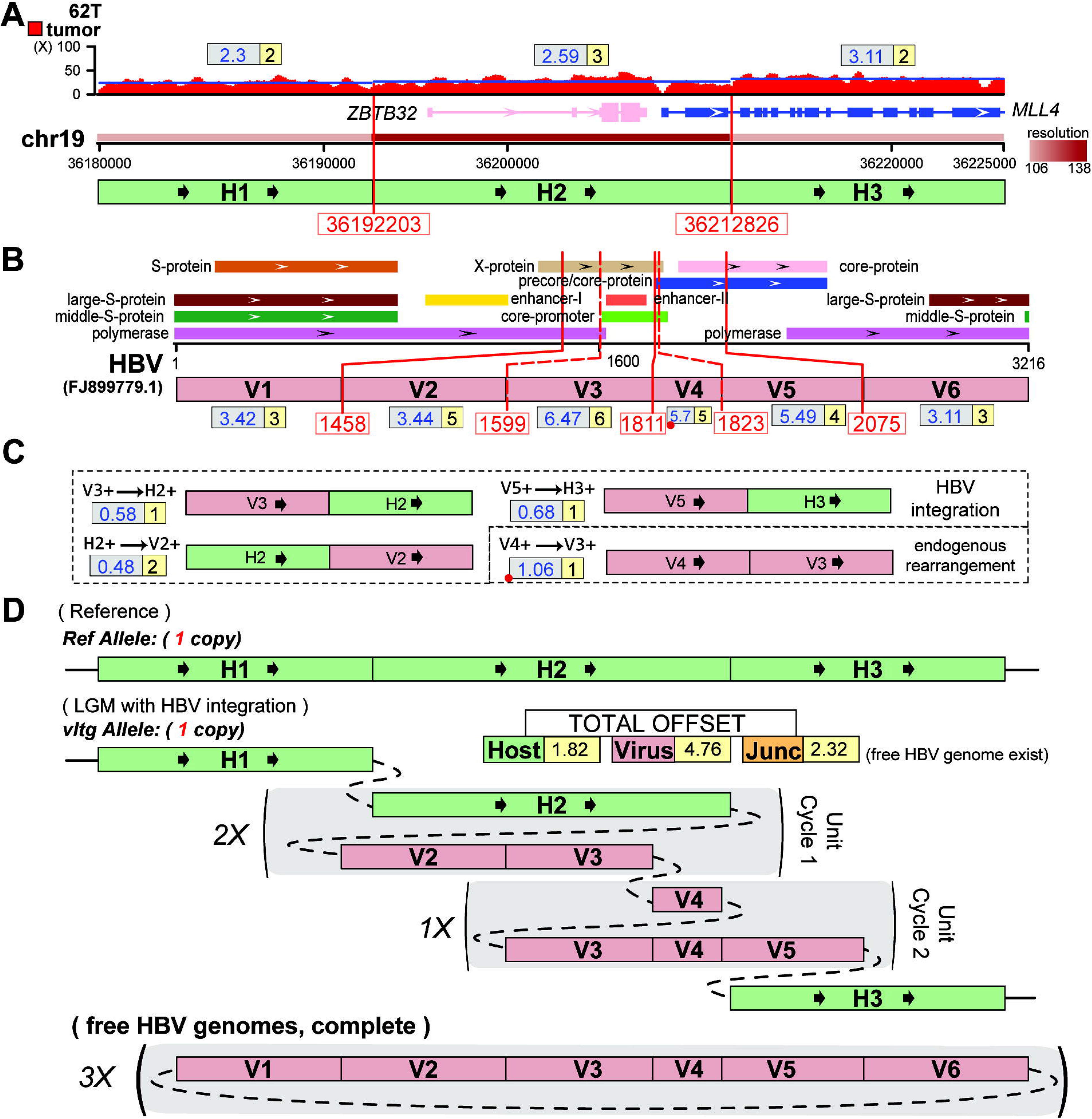
Presentative LGM at HBV integration sites (gene MLL4) on chr 19 (minor allele) of the 62T HCC sample [11]. (**A**) Human genomic region flanking HBV integrations are divided into three segments (H1 ~ H3) by VITs denoted with breakpoints. Depth spectrum is displayed with original (grey frame) and ILP-adjusted (yellow frame) copy numbers of segments. (**B**) Segmentation (V1 ~ V6) of HBV genome by VITs and SVs. The segment less than 100bp is marked with red dot. (**C**) Variant segment junctions utilized in Conjugate graph. (**D**) Resolved alleles of the ‘Simplest LGM’ are indicated as string of coloured segments with copy times, including reference allele and that harbours HBV integrations. Sum of absolute offset of segments and junctions are shown. Unit-cycles (shadowed areas) in LGM are denoted with repeat time. Note that the free HBV genomes might exist.

There are two HBV-integrated LGMs located on the heterozygous diploid (62T and 182T, **Supplementary Figure S12A-B**). The sample 62T has three HBV integrations at the MLL4 oncogene on chr19q13.12 (**Figure 3**). The LGM contains two unit-cycles. The NO.1 unit-cycle duplicates the ‘H2’ host segment covering the whole gene body of ZBTB32, the promoter and first two exons of MLL4, which are abundant of the regulatory element marks [23]. The structure is predicted to have no truncation effect on gene MLL4 as the coding frame is completely preserved (‘H2’ to HBV to ‘H3’). Furthermore, the inserted HBV segments (‘V2’ to ‘V5’) carry the enhancer II and core promoter [24, 25], which potentially promote the MLL4 expression [11]. The HBV DNA segments inserted at gene NBAS on chr2p24.3 of sample 182T, and resulted in the deletion of the host segment (‘H2’) containing 14 exons of NBAS (**Supplementary Figure S13**). The resolved LGM has a simple structure that the ‘H2’ host segment is replaced by the majority of the HBV genome.

We found that the sample 13T and 260T both have copy number gains of broad regions at the HBV integration loci. The 13T has HBV integrations at the TERT gene locus on chr5p15.33, which is heterozygous triploid extending to p15.2 (**Supplementary Figure S12C**). As the second host segment (‘H2’, CN = 2, intron region) has one copy loss, we determined that the HBV integrated on the minor allele and replaced the ‘H2’ segment (**Supplementary Figure S14**). The HBV genome is divided into 16 segments by breakpoints of two integrations and seven viral rearrangements, which are all considered in the LGM construction. The inserted HBV DNA segments formed a complex structure consisting of four unit-cycles. The unit-cycle NO.2 and NO.3 carry the HBV enhancers and core promoters, which might explain the elevated gene expression of TERT in this sample [11]. In sample 260T, the HBV integrated at the ANK3 gene on chr10 (heterozygous tetraploid, minor CN = 2, **Supplementary Figure S12D**). The resolved LGM contains two unit-cycles including two HBV integrations and three HBV SVs (**Supplementary Figure S15**). These two unit-cycles are completely composed of inserted HBV DNA segments and substitute the ‘H2’ host segment in two allele copies. Notably, our algorithm reports the unit-cycle NO.1 only in one allele, and it might exist in the form of ecDNA.

The remaining two HCC samples displayed arm-level LOH of HBV-integrated chromosomes (101T and 261T). The sample 101T has two HBV integrations at the GLRA2 gene locus in the chrXp22.2 (CN = 1). The remained heterozygous status of the chrXq supports the chrXp is an LOH (**Supplementary Figure S12E**). The LGM is simple and the only unit-cycle is the inserted ‘V2’ HBV segment that replaces the ‘H2’ host segment (partial intron of GLRA2, **Supplementary Figure S16**). In sample 261T, there are two HBV integrations located in the intron of BBS2 gene on chr16 (CN = 1, **Supplementary Figure S12F**). Similar to the sample 13T, the HBV genome is divided into 17 segments by two integrations and seven HBV SVs. Our algorithm reports the LGM consisting of three unit-cycles with all segments from the viral genome (**Supplementary Figure S17**). These unit-cycles have complex structures composed of deletion, tandem duplication, and inversions. Again, one host deletion (‘H2’) is caused by the inserted HBV DNA.

### Diverse affection patterns of oncovirus integrations

According to the resolved LGMs, we summarized five affection patterns of the oncovirus integrations in our samples. Firstly, the integrations locate in a single intron, including the HPV16 in T4931 (gene GLI2), and the HBV in 13T (TERT), 101T (GLRA2), 260T (ANK3), and 261T (BBC2). The inserted viral DNA segments carry the epigenomic regulator elements, such as the long control region (LCR) of HPV, and the enhancers and core promoter of HBV, which might contribute to the local expression dysregulations. The abnormal alternative splicings are also reported in researches of both HPV and HBV integrations [2, 13]. Secondly, the LGM covers multiple exons in sample UM-SCC-47 (HPV16 at TP63) and 182T (HBV at NBAS), both likely lead to the truncation of the coding-frame. Thirdly, in sample UPCI-SCC090, the LGM contains focal hyper-amplifications harboring the complete genes (PIM1 and SNORD112), which commonly relate to elevated expression and might induce tumorigenesis [26]. Fourthly, in sample 62T, tandem duplication doubles the promoter and maintains the complete coding-frame of MLL4, which is also possible subject to the inserted HBV enhancer and core promoter. Fifthly, both of T6050 sample and SiHa cell line have HPV integrations at the intergenic regions between gene KLF5 and KLF12, whose expressions are probably affected by the amplified HPV LCR in the LGM. The HeLa cell line has been reported that the inserted HPV segments have long-range chromatin interactions with downstream MYC gene, which has high RNA expression phased in the HPV-integrated haplotype [1]. This implicates the cis-regulation of HPV integration is possibly mediated by the epithelium specific viral enhancer remained in the LGMs.

### Enhanced Microhomology-mediated mechanism of HPV and HBV integration

The resolved LGMs enable us to analyze the details of breakpoints and investigate the oncovirus integration mechanisms. Here, we propose that the slipped microhomology and the small junctional insertion (see **Methods**), which are the defining characteristic of the microhomology-mediated end joining [27, 28]. A previous study on cervical cancer has reported the enrichment of aligned microhomology at HPV integration sites [7]. In total, 54.05% (20/37) of virus integrations have aligned microhomology at the junction site. Meanwhile, the percentage became to 83.78% (31/37) when the slipped microhomology and the junctional insertion are considered (**Figure 4, Supplementary Table S8, Supplementary Figure S19-S26**), and it is more prevalent in HPV integrations (91.67%, 22/24) comparing with HBV (69.23%, 9/13). Moreover, microhomology and insertions are found at the junction sites of the majority (84.84%, 28/33) of the endogenous SVs: 100% (9/9) for host SVs in HPV LGMs, 83.33% (5/6) for HPV SVs, and 77.78% (14/18) for HBV SVs. Furthermore, the complex SVs in sample GBM0152 also shows similar phenomenon that 60% (12/20) of the SV junctions carry the microhomology (**Supplementary Table S7** and **S8, Supplementary Figure S18, S27** and **S28**). Microhomology enrichment supports the suggested replication-based mechanism, which generates these complex rearrangements in this sample [15].

**Figure 4.**
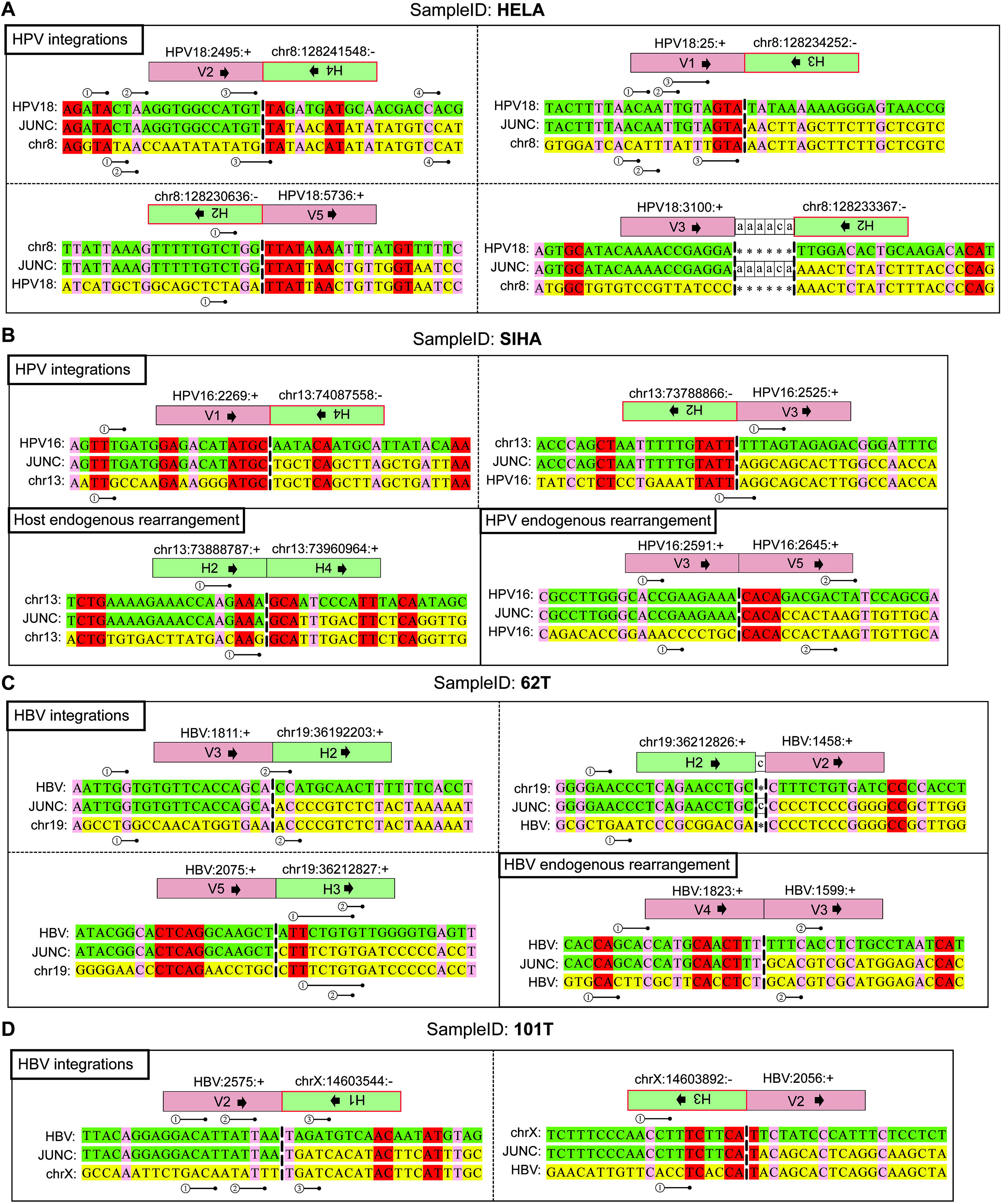
Alignment of the sequence around the integration site between the host and virus genome, and the endogenous rearrangements. In the samples (**A**) HeLa, (**B**) SiHa, (**C**) 62T, (**D**) 101T. The junction boundaries are shown as vertical dashed lines. All viral sequences are from the reference strand. Green, upstream junction partner; yellow, downstream junction partner; red, nucleotides that vertically align to both reference sequences (aligned microhomologous bases); pairwise numbered sticks, slipped microhomologous bases. All microhomologous bases follow the 5’-to-3’ direction. The junction segment IDs are corresponding to the segments in the resolved LGMs.

We further propose that the enhanced microhomology-mediated end joining (MMEJ) may be the underlying mechanism of both HPV and HBV integrations together with the endogenous host and viral rearrangements in the local region. Each unit-cycle in the resolved LGMs has at least one boundary harboring the microhomology or junctional insertion (**Figure 5** for HPV16, **Supplementary Figure S29** for HPV18, and **Figure 6** for HBV), suggesting that the formation of each unit-cycle is possibly initialized via the replication-based template-switching event. The well-known alternative mechanisms of MMEJ include the fork stalling and template switching (FoSTeS) [29] and the microhomology-mediated break-induced replication (MMBIR) [30].

**Figure 5.**
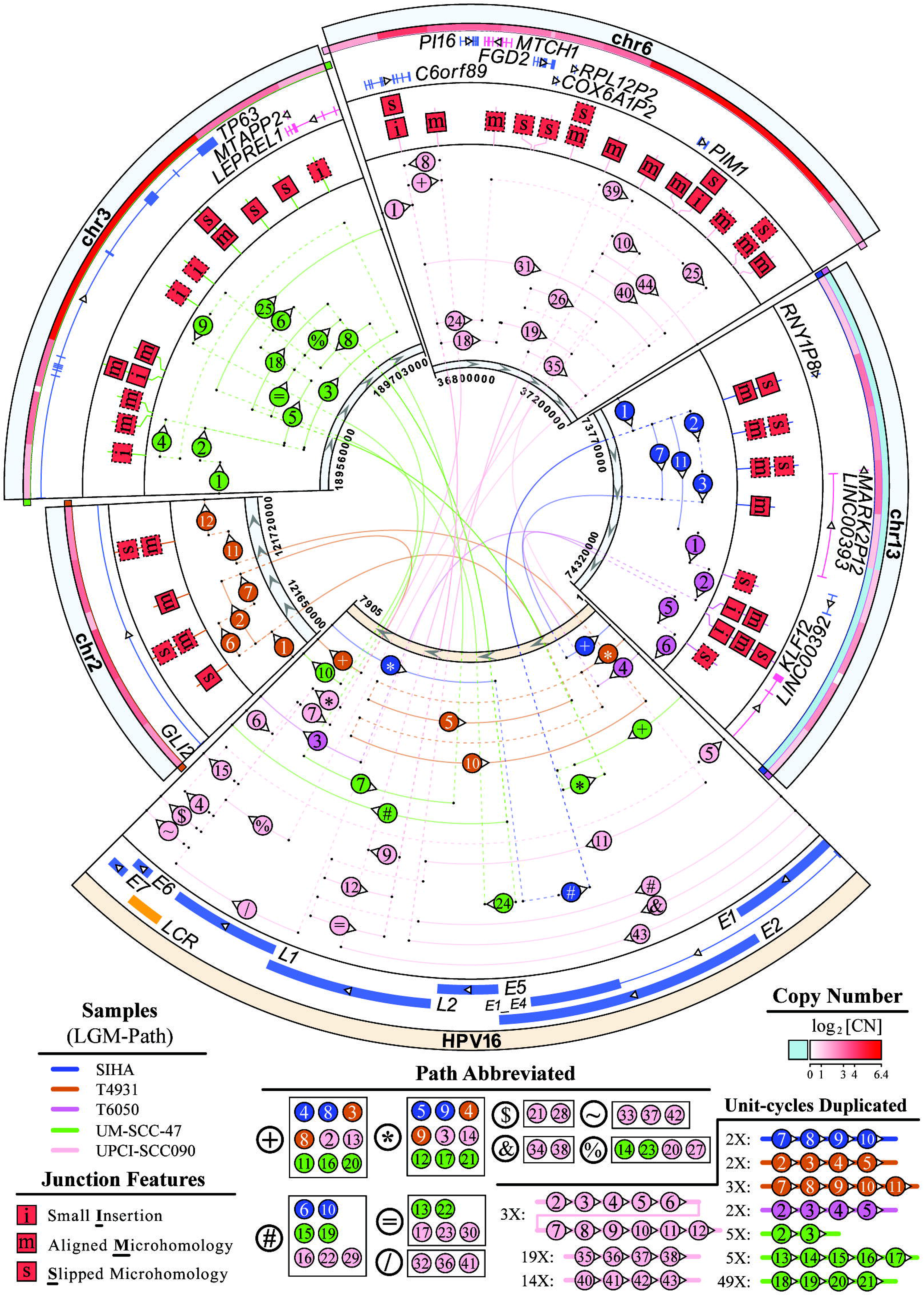
Features of HPV16 integrated LGMs in five samples. Human genomic segments related to HPV16 integrated LGMs are shown as sectors with their relevant LGM path in sample specific colours. Segments of LGMs are denoted by numbered concentric arcs. The numbers, starting from 1, indicate the order in which each arc is visited starting from the source segment of an LGM. Each special symbol shown in “Path Abbreviated” section in the figure legends denotes an arc that is visited more than once in the LGM, such as the ‘+’ symbol for NO.4 and NO.8 arcs in LGM of SiHa sample. Sequential segments in LGM could be merged in single arc (**Supplementary Table S9**). Repeat times of unit-cycles in LGMs are stated in figure legend. Features of HPV16 integration (solid box) and host rearrangement (dotted) sites are depicted as single-letter icons. DNA copy number (CN) is displayed in gradient red colour (light-blue for regions outside of the LGM), with bilateral labels in relevant sample colour. The HPV16 genome reference is NC 001526.4 from the NCBI Nucleotide database.

**Figure 6.**
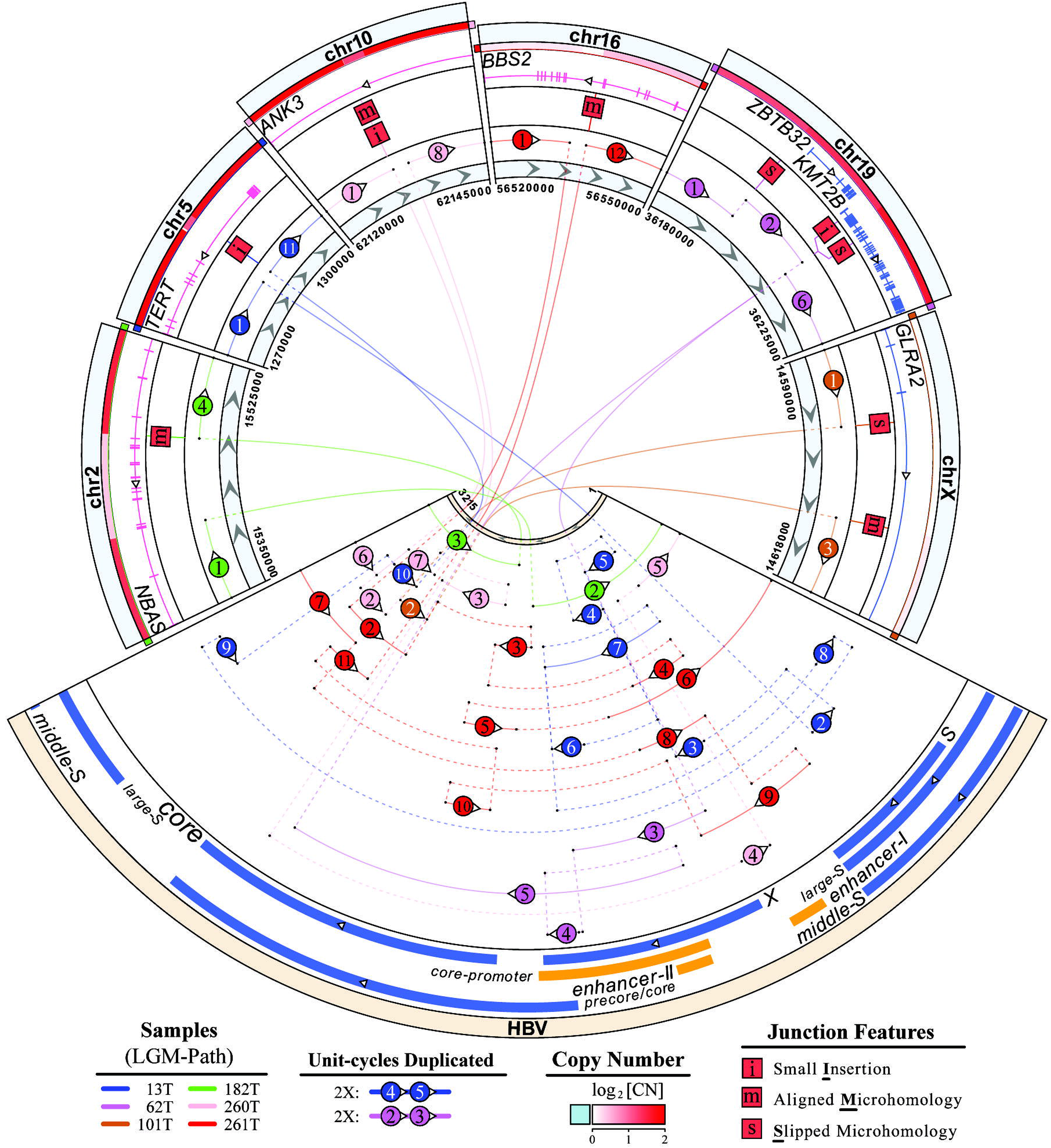
Features of HBV integrated LGMs in six HCC samples. Human genomic segments related to HBV integrated LGMs are shown as sectors with their relevant LGM path in sample specific colours. Segments of LGMs are denoted by numbered concentric arcs. The numbers, starting from 1, indicate the order in which each arc is visited starting from the source segment of an LGM. Sequential segments in LGM could be merged in single arc (**Supplementary Table S9**). Repeat times of unit-cycles in LGMs are stated in figure legend. Features of HBV integrated sites are depicted as single-letter icons. DNA copy number (CN) is displayed in gradient red colour (light-blue for regions outside of the LGM), with bilateral labels in relevant sample colour. The HBV genome reference is FJ899779.1 from the NCBI Nucleotide database.

We hypothesize that the HPV integrations might mainly occur during the DNA replication process. All of the HPV-integrated LGMs harbor the focal duplications of human genomic segments (**Figure 5** and **Supplementary Figure S29**), while most (83%, 5/6) of the HBV-integrated LGMs only contain focal deletions caused by the substitution of the inserted viral segments (**Figure 6**). Additionally, most (83%, 5/6) of the HPV-integrated LGMs have host SVs, but none of the HBV-integrated LGMs has. These differences further enlightens us to hypothesize that the HPV integrations might mainly occur during the DNA replication process (i.e., the S phase during the mitosis) and form complex structure via the rolling circle replication [2, 31], and the HBV integrations likely arise under the DNA repairing circumstance and are prone to produce simple structure.

## Discussion

The detection of the oncovirus integrations and structural variations is widely developed, and the effects and formation mechanisms of integrations are extensively studied. However, the investigation in the local genomic region with the linear complex structure remains limited. Our conjugate graph algorithm constructs the local genomic maps of HPV and HBV integrations. It provides details to research the regulatory influences on local genes and the underlying formation mechanisms, combining with the copy changes of rearranged host and viral genomic segments.

The local genomic map sometimes contains segments shorter than the insert-size of the paired-end sequencing data or harboring repeated DNA sequence, leading to the underestimated copy number calculation from the average depth of the relevant segment. Similarly, the insufficient split-reads count also reduces the copy number of the SV and VIT junctions. These inaccurate copy numbers will hinder the accuracy of the LGM results. The integer linear programming in our algorithm overcomes such difficulties by assigning smaller weights for relevant vertex and edge in the graph model, and successfully construct the optimal LGM with minimum offsets.

The unit-cycles in the conjugate graph model are the basic structural units representing the connections of segments and junctions. Unit-cycles are merged in ways to form different LGM solutions. Due to the short reads in NGS data, the factual orders of merging cannot be determined technically. Here, we select the solution with the simplest structure to report, following Ocham’s razor. The candidate LGMs could be further filtered based on the phased structure obtained from the advanced DNA sequencing, such as 10x long-range linked-reads and single-molecule reads. Furthermore, when the virus-integrated chromosome is heterozygous unbalanced polyploidy, it is difficult to determine whether the paternal or maternal allele carries the viral insertions based on short reads. The mutations phasing between the local genomic segments and the flanking regions of the constructed LGMs will help solve the problem. Multiple sequencing technology can provide this additional information, including the mate-pair library with large insert-size [1], 10x linked-reads [32], and Hi-C [33].

When the virus-integrated allele has more than one copy, our algorithm will try to distribute all relevant unit-cycles evenly among the multiple copies to minimize the differences. However, the actual count of the unit-cycles might change within different allele copies (e.g., **Supplementary Figure S5D** and **S6D**). According to the circular structure and common hyperamplification, the unit-cycles can be the circular extrachromosomal DNA, which needs more investigations in the future study, for example, the inter-chromosomal interactions from Hi-C data [34]. Recent studies have proved that the ecDNA promotes oncogene expressions and drives tumor evolution [35, 36]. The speculated oncovirus integrations related ecDNAs probably have similar carcinogenic functions.

From the constructed LGMs, we can get the structural investigation on the formation mechanism of the oncovirus integrations and endogenous SVs in the local genomic region. To our knowledge, this is the first time to combine the aligned/slipped microhomology and junctional insertion together in the mechanism research of viral integrations. The enrichment of microhomology and insertion indicates that both the HPV and HBV integrations may be generated via the microhomology-mediated end joining (MMEJ), promoted during the HPV16 infection [37]. Furthermore, the focal duplicates and host SVs are frequent in the HPV-integrated LGMs, while the HBV-integrated LGMs are abundant in focal deletions and complex virus SVs. This apparent difference suggests that the formation circumstances of HPV and HBV integration may be different. To answer this question, it needs more samples and experiments in a future study.

In summary, we present a conjugate graph algorithm to construct the local genomic map of virus integrations mediated host and viral rearrangements from the whole genome sequencing data. The algorithm can normalize imperfect sequencing depths and elucidate the linear DNA structure at virus integration sites. The simulation tests and the applications on WGS data of cancer samples prove the reliability and credibility of our model. The results shed light on the genomic dysregulation and formation mechanism of the oncovirus integrations.

## Data availability

The conjugate graph algorithm code, LGM analysis data of all samples (Supplementary Dataset zip file), and LGM construction pipeline demo are available in github repository (https://github.com/deepomicslab/FuseSV). The cancer whole genome sequencing data is obtained under SRA accession number SRA180295 [7], and EGA accession number EGAS00001000599 [2] and ERP001196 [11].

## Author contributions

S.L. proposed the conjugate graph model and designed algorithm to obtain the LGM paths. W.J. and C.X. implemented and optimized the algorithms. W.J. and C.X. performed the WGS data analysis. C.X. performed the simulation work. W.J. developed the FuseSV. W.J. summarised and compared the viral integration features and mechanisms. W.J., C.X. and S.L. wrote and revised the manuscript. All authors reviewed the article and approved the final manuscript.

## Author description

Wenlong Jia received his Ph.D. degree from the Department of Computer Science of City University of Hong Kong in 2019. His research interests include cancer genetics, bioinformatics, and computational biology.

Chang Xu received his master degree of computational data science from Carnegie Mellon University in 2019. His research interests include bioinformatics, algorithm development, and data mining.

Shuai Cheng Li is currently a professor and head of a lab at the Department of Computer Science of City University of Hong Kong. The Li lab develops various computational tools to gain insight into the interactions between omics data and disease phenotypes. Her research interests include genetics, bioinformatics, and machine learning.

## Supplementary Data

**Supplementary Table S1.** Details of simulated oncovirus integrated local genomic map.

**Supplementary Table S2.** Mutations used to reconstruct HPV genomes in six HPV-infected cancer samples.

**Supplementary Table S3.** Details of local genomic map in four cervical carcinoma samples.

**Supplementary Table S4.** Details of local genomic map in two HPV infected HNSCC cancer cell lines.

**Supplementary Table S5.** Mutations used to reconstruct HBV genomes in six HBV-infected HCC samples.

**Supplementary Table S6.** Details of local genomic map in six HBV infected HCC samples.

**Supplementary Table S7.** Local genomic map of one complex SV on chr12 of GBM0152 sample.

**Supplementary Table S8.** MHs and insertions at the junction sites in resolved local genomic maps.

**Supplementary Table S9.** Relationship of segment IDs in LGMcircos and LGM.

**Supplementary Figure S1.** The eight LGM modes in simulation work are denoted with features and represented by basic structures.

**Supplementary Figure S2.** Copy number distribution along the HPV-integrated chromosome in four cervical cancer samples [7].

**Supplementary Figure S3.** Presentative LGM at HPV integration sites (gene GLI2) on chr2 (major allele) of the T4931 sample [7].

**Supplementary Figure S4.** Presentative LGM at HPV integration sites (upstream of gene MYC) on chr8 (major allele) of the HeLa cell line [7].

**Supplementary Figure S5.** Presentative LGM at HPV integration sites (gene KLF12) on chr13 (LOH) of the T6050 sample [7].

**Supplementary Figure S6.** Presentative LGM at HPV integration sites (upstream of gene KLF12) on chr13 (LOH) of the SiHa cell line [7].

**Supplementary Figure S7.** Copy number distribution along the HPV-integrated chromosome in two HNSCC cell lines [2].

**Supplementary Figure S8.** Presentative LGM at HPV integration sites (gene TP63) on chr3 (minor allele) of the UM-SCC-47 cell line [2].

**Supplementary Figure S9.** Presentative LGM at HPV integration sites (gene TP63) on chr3 (major allele) of the UM-SCC-47 cell line [2].

**Supplementary Figure S10.** Presentative LGM at HPV integration sites (gene PIM1) on chr6 (minor allele) on chr6 of the UPCI-SCC090 cell line [2].

**Supplementary Figure S11.** Presentative LGM at HPV integration sites (gene PIM1) on chr6 (major allele) of the UPCI-SCC090 cell line [2].

**Supplementary Figure S12.** Copy number distribution along the HBV-integrated chromosome in six HCC samples [11].

**Supplementary Figure S13.** Presentative LGM at HBV integration sites (gene NBAS) on chr2 of the 182T HCC sample [11].

**Supplementary Figure S14.** Presentative LGM at HBV integration sites (gene TERT) on chr5 (minor allele) of the 13T HCC sample [11].

**Supplementary Figure S15.** Presentative LGM at HBV integration sites (gene ANK3) on chr10 of the 260T HCC sample [11].

**Supplementary Figure S16.** Presentative LGM at HBV integration sites (gene GLRA2) on chrX of the 101T HCC sample [11].

**Supplementary Figure S17.** Presentative LGM at HBV integration sites (gene BBS2) on chr16 of the 261T HCC sample [11].

**Supplementary Figure S18.** Presentative LGM at complex SV locus on chr12 of the GBM0152 cancer sample [15].

**Supplementary Figure S19.** Alignment of the sequence around the integration site between the human genome and the HPV16 genome, and the endogenous rearrangements on human genome and the HPV16 genome respectively, in the T4931 sample [7].

**Supplementary Figure S20.** Alignment of the sequence around the integration site between the human genome and the HPV16 genome, and the endogenous rearrangements on human genome, in the T6050 sample [7].

**Supplementary Figure S21.** Alignment of the sequence around the integration site between the human genome and the HPV16 genome, and the endogenous rearrangements on human genome and the HPV16 genome respectively, in the UM-SCC-47 cell line [2].

**Supplementary Figure S22.** Alignment of the sequence around the integration site between the human genome and the HPV16 genome, and the endogenous rearrangements on human genome and the HPV16 genome respectively, in the UPCI-SCC090 cell line [2].

**Supplementary Figure S23.** Alignment of the sequence around the integration site between the human genome and the HBV genome, and the endogenous rearrangements on HBV genome, in the 13T HCC sample [11].

**Supplementary Figure S24.** Alignment of the sequence around the integration site between the human genome and the HBV genome in the 182T HCC sample [11].

**Supplementary Figure S25.** Alignment of the sequence around the integration site between the human genome and the HBV genome, and the endogenous rearrangements on HBV genome, in the 260T HCC sample [11].

**Supplementary Figure S26.** Alignment of the sequence around the integration site between the human genome and the HBV genome, and the endogenous rearrangements on HBV genome, in the 261T HCC sample [11].

**Supplementary Figure S27.** Alignment of the sequence around the endogenous complex rearrangements on human genome in the GBM0152 cancer sample [15].

**Supplementary Figure S28.** Features of the complex SV LGM in GBM0152 sample. Human genomic segments related to complex SVs LGMs are shown as sectors with their relevant LGM path.

**Supplementary Figure S29.** Features of HPV18 integrated LGM in HeLa cell line.

