## Supplementary Figures for "Resolving complex structures at oncovirus integration loci with conjugate graph"

* To whom correspondence should be addressed.

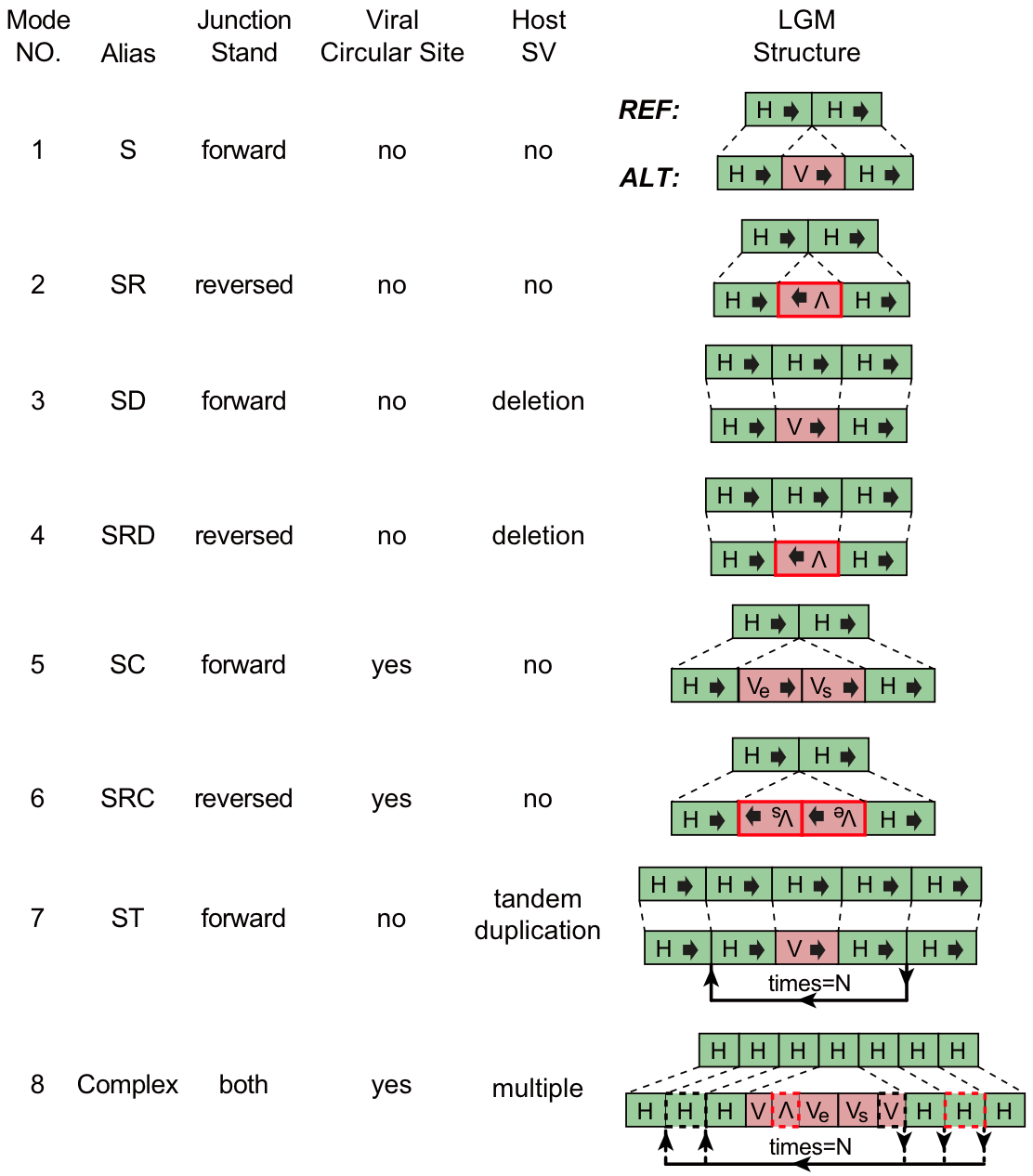

**Supplementary Figure S1.** **The eight LGM modes in simulation work are denoted with features and represented by basic structures**. The alteration sequences with host (‘H’) and virus (‘V’) segments are aligned with the reference. Segments in red frame are reverse complementary counterpart. The first and last viral segments are denoted by ‘V_s_’ and ‘V_e_’, respectively. Dashed frame and directed lines means segments and paths randomly selected.

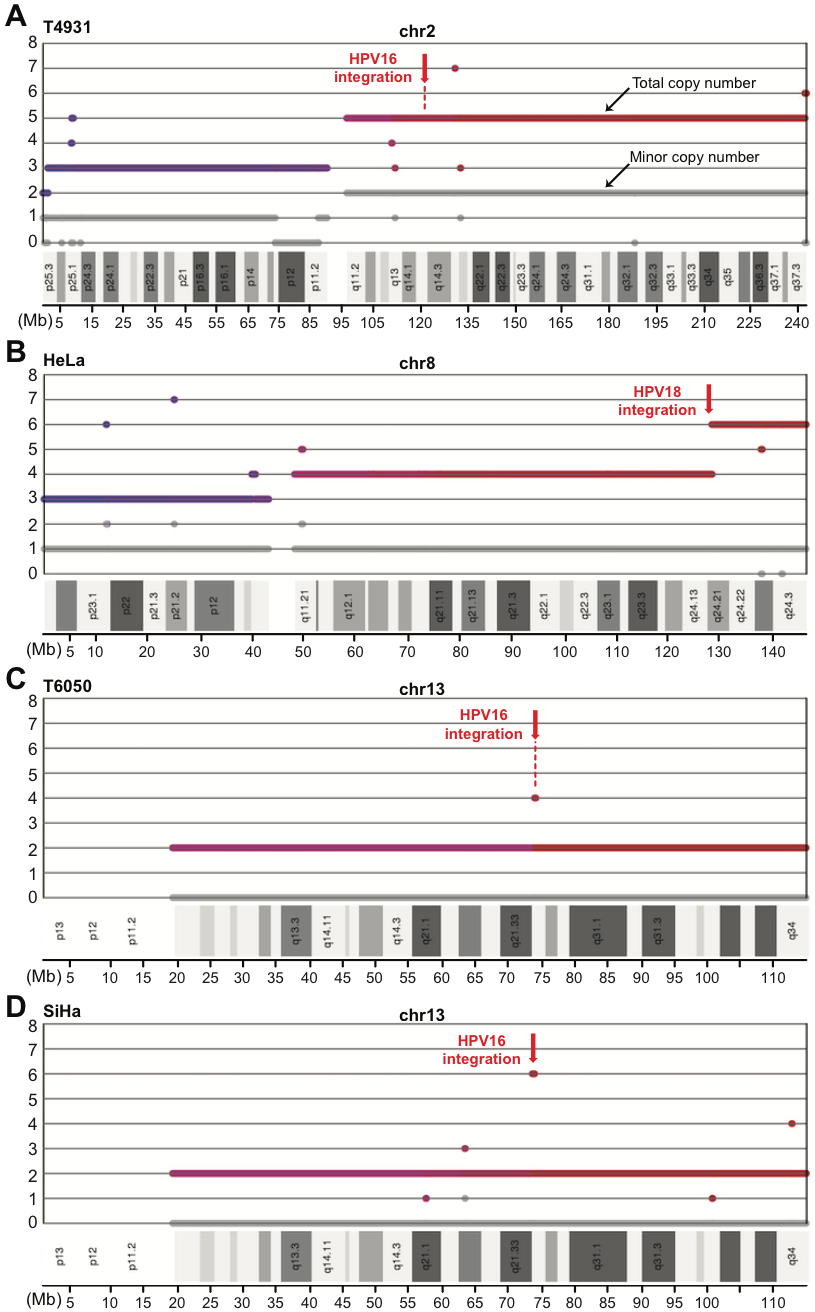

**Supplementary Figure S2.** **Copy number distribution along the HPV-integrated chromosome in four cervical cancer samples** [7]. Figures are from patchwork results. HPV integration sites are denoted by red arrows. Colored and grey bold lines indicate total and minor copy number respectively.

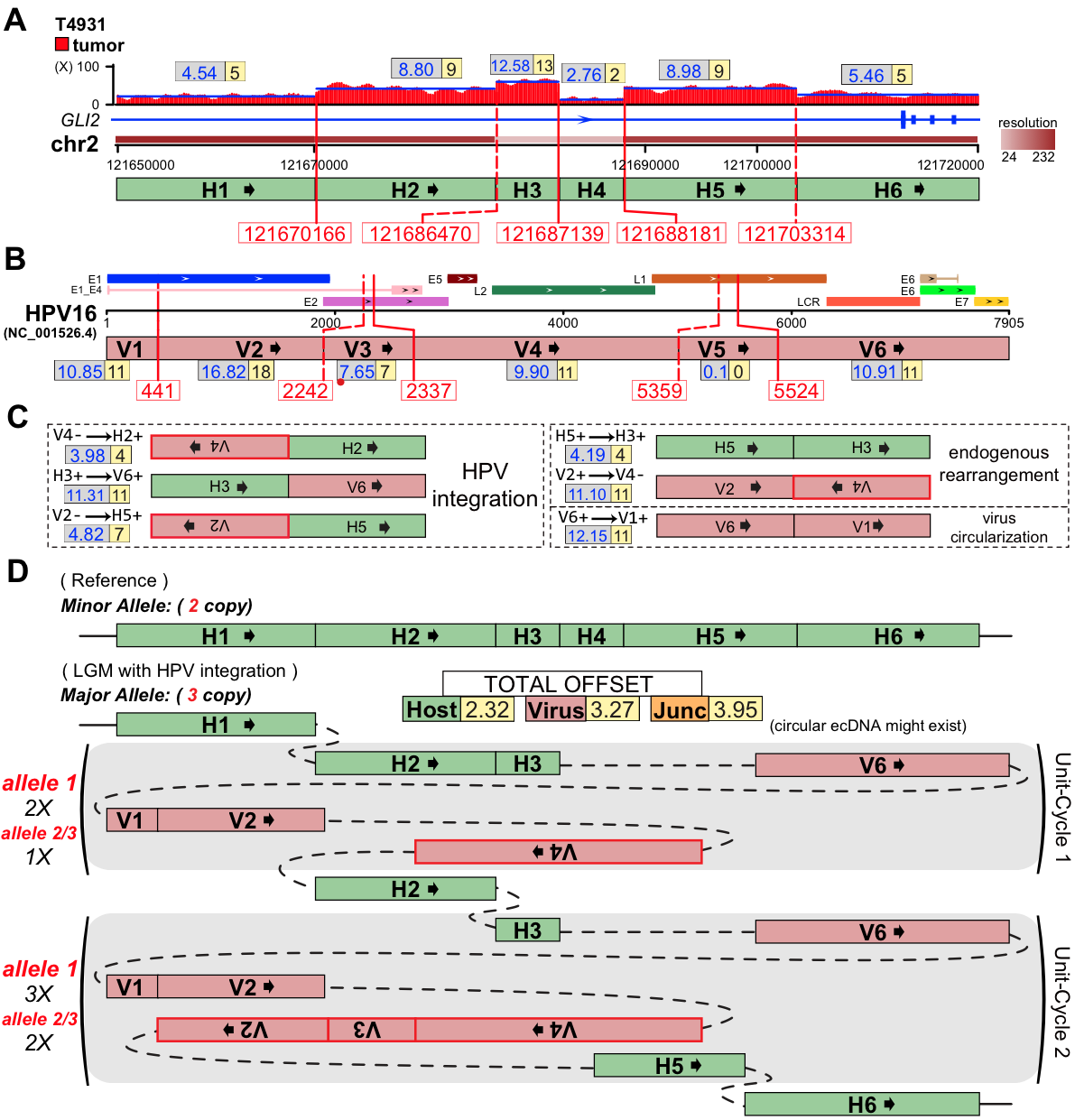

**Supplementary Figure S3.** **Presentative LGM at HPV integration sites (gene *GLI2*) on chr2 (major allele) of the T4931 sample** [7]. (**A**) Human genomic region flanking HPV integrations are divided into six segments (H1∼H6) by VITs (red solid-line) and SVs (red dashed-line) denoted with breakpoints. Depth spectrum is displayed with original (grey frame) and ILP-adjusted (yellow frame) copy numbers of segments. (**B**) Segmentation (V1∼V6) of HPV16 genome by VITs and SVs. The segment (V3) less than 100bp is marked with red dot. (**C**) Variant segment junctions utilized in Conjugate graph. (**D**) Resolved alleles of the ‘Simplest LGM’ are indicated as string of coloured segments with copy times, including reference allele (minor allele) and that harbours HPV16 integrations (major allele). Sum of absolute offset of segments and junctions are shown. Unit-cycles (shadowed areas) in LGM are denoted with repeat time. Note that the HPV-integrated alleles might have different copies of unit-cycles, which might exist as circular ecDNAs.

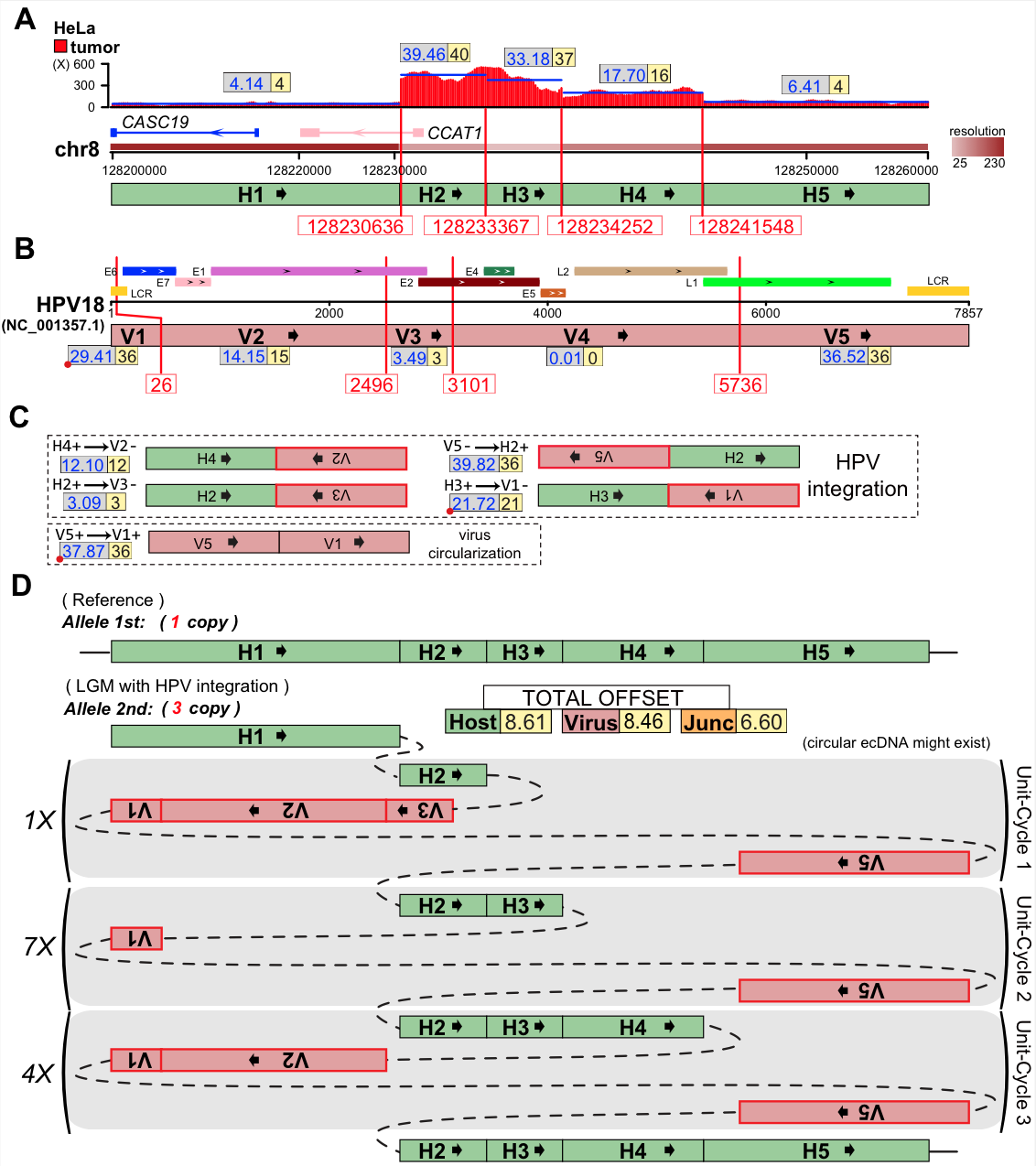

**Supplementary Figure S4.** **Presentative LGM at HPV integration sites (upstream of gene *MYC*) on chr8 (major allele) of the HeLa cell line** [7]. (**A**) Human genomic region flanking HPV integrations are divided into five segments (H1∼H5) by VITs denoted with breakpoints. Depth spectrum is displayed with original (grey frame) and ILP-adjusted (yellow frame) copy numbers of segments. (**B**) Segmentation (V1∼V5) of HPV18 genome by VITs. The segment (V1) less than 100bp is marked with red dot. (**C**) Variant segment junctions utilized in Conjugate graph. (**D**) Resolved alleles of the ‘Simplest LGM’ are indicated as string of coloured segments with copy times, including reference allele (minor allele) and that harbours HPV18 integrations (major allele). Sum of absolute offset of segments and junctions are shown. Unit-cycles (shadowed areas) in LGM are denoted with repeat time. The unit-cycles might exist in the form of circular ecDNA.

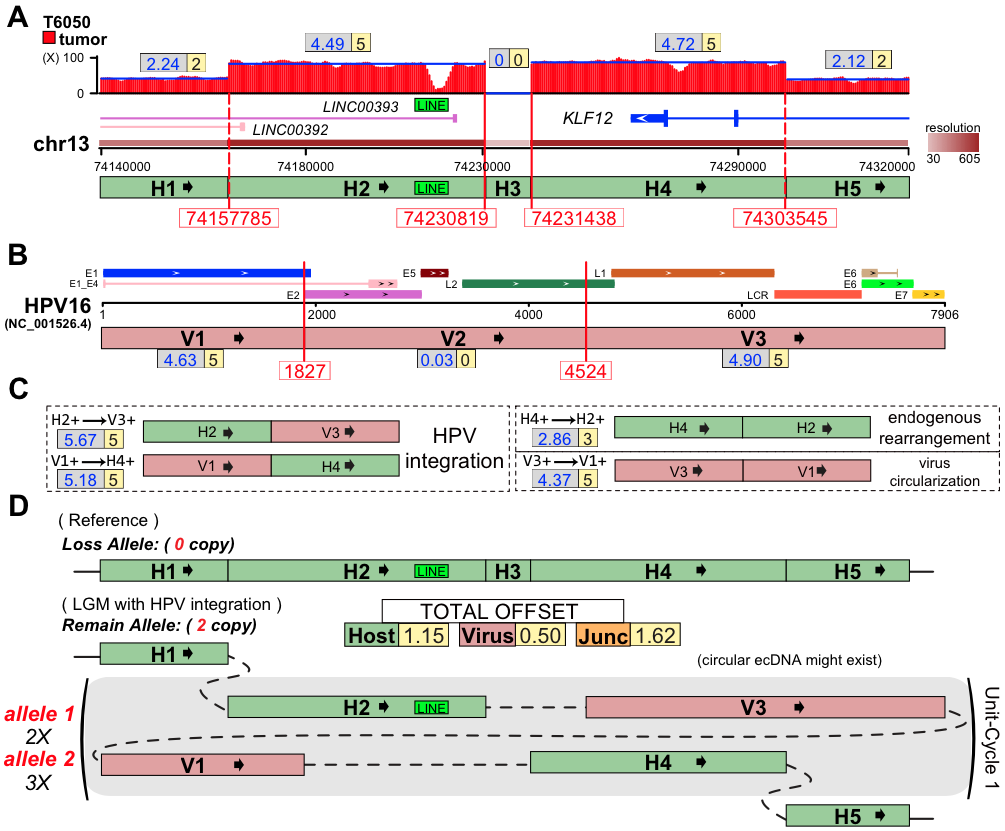

**Supplementary Figure S5.** **Presentative LGM at HPV integration sites (gene *KLF12*) on chr13 (LOH) of the T6050 sample** [7]. (**A**) Human genomic region flanking HPV integrations are divided into five segments (H1∼H5) by VITs (red solid-line) and SVs (red dashed-line) denoted with breakpoints. Depth spectrum is displayed with original (grey frame) and ILP-adjusted (yellow frame) copy numbers of segments. (**B**) Segmentation (V1∼V3) of HPV16 genome by VITs and SVs. (**C**) Variant segment junctions utilized in Conjugate graph. (**D**) Resolved alleles of the ‘Simplest LGM’ are indicated as string of coloured segments with copy times, including reference allele (loss allele) and that harbours HPV16 integrations (remain allele). Sum of absolute offset of segments and junctions are shown. Unit-cycles (shadowed areas) in LGM are denoted with repeat time. Note that the HPV-integrated alleles might have different copies of unit-cycles, which might exist as circular ecDNAs.

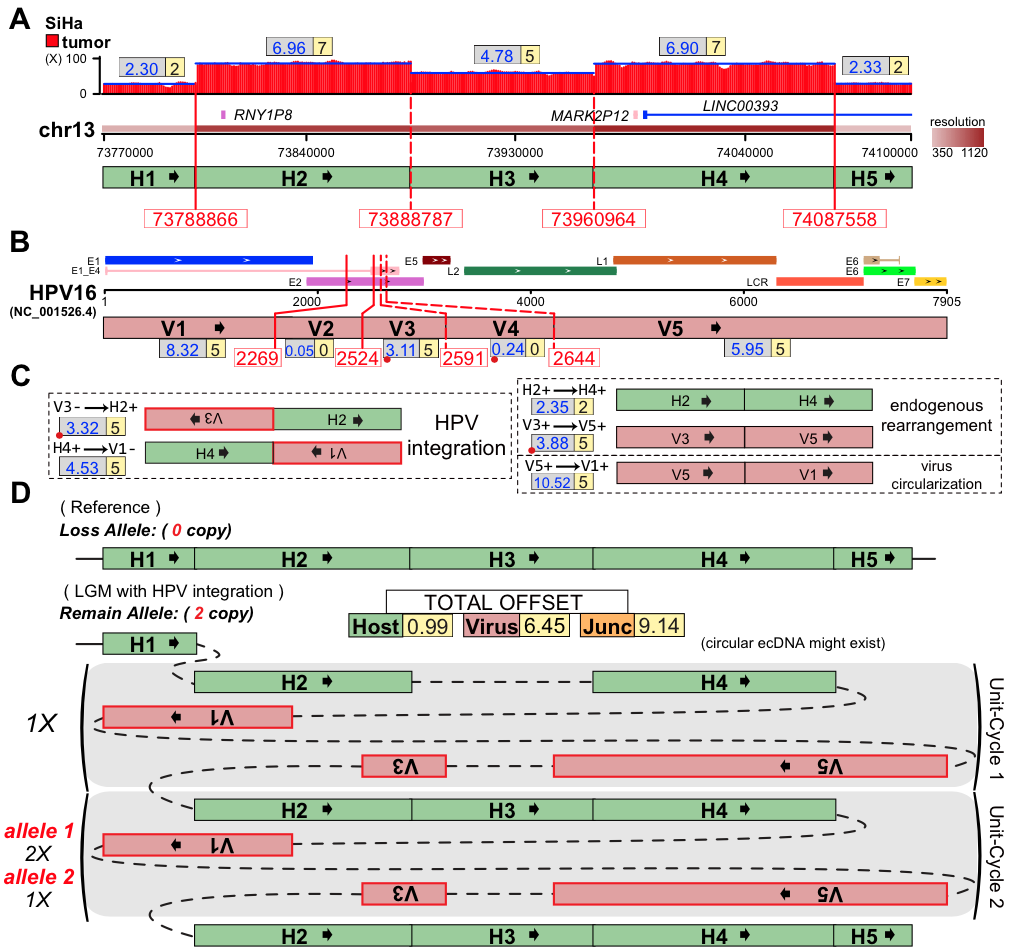

**Supplementary Figure S6.** **Presentative LGM at HPV integration sites (upstream of gene *KLF12*) on chr13 (LOH) of the SiHa cell line** [7]. (**A**) Human genomic region flanking HPV integrations are divided into five segments (H1∼H5) by VITs (red solid-line) and SVs (red dashed-line) denoted with breakpoints. Depth spectrum is displayed with original (grey frame) and ILP-adjusted (yellow frame) copy numbers of segments. (**B**) Segmentation (V1∼V5) of HPV16 genome by VITs and SVs. (**C**) Variant segment junctions utilized in Conjugate graph. (**D**) Resolved alleles of the ‘Simplest LGM’ are indicated as string of coloured segments with copy times, including reference allele (loss allele) and that harbours HPV16 integrations (remain allele). Sum of absolute offset of segments and junctions are shown. Unit-cycles (shadowed areas) in LGM are denoted with repeat time. Note that the HPV-integrated alleles might have different copies of unit-cycles, which might exist as circular ecDNAs.

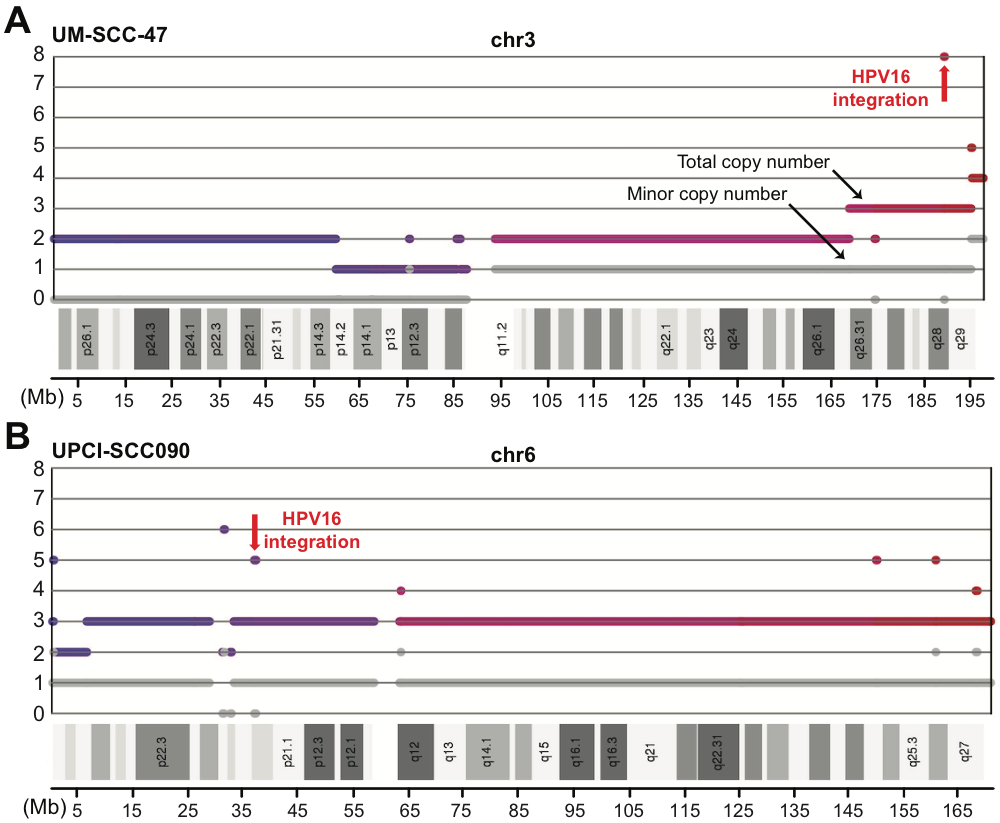

**Supplementary Figure S7.** Copy number distribution along the HPV-integrated chromosome in two HNSCC cell lines [2]. Figures are from patchwork results. HPV integration sites are denoted by red arrows. Colored and grey bold lines indicate total and minor copy number respectively.

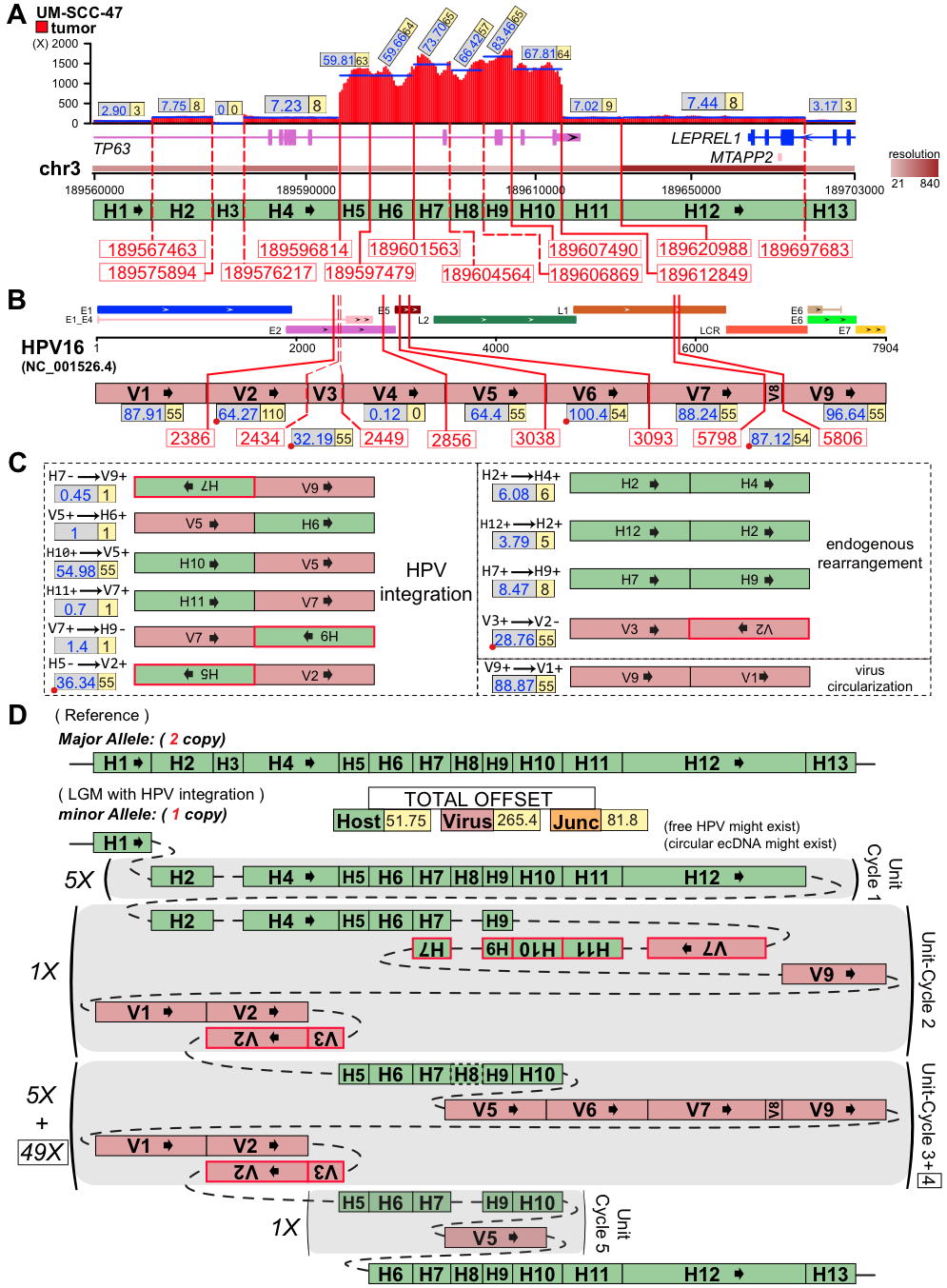

**Supplementary Figure S8.** **Presentative LGM at HPV integration sites (gene *TP63*) on chr3 (minor allele) of the UM-SCC-47 cell line** [2]. (**A**) Human genomic region flanking HPV integrations are divided into 13 segments (H1∼H13) by VITs (red solid-line) and SVs (red dashed-line) denoted with breakpoints. Depth spectrum is displayed with original (grey frame) and ILP-adjusted (yellow frame) copy numbers of segments. (**B**) Segmentation (V1∼V9) of HPV16 genome by VITs and SVs. The segments less than 100bp are marked with red dot. (**C**) Variant segment junctions utilized in Conjugate graph. (**D**) Resolved alleles of the ‘Simplest LGM’ are indicated as string of coloured segments with copy times, including reference allele (major allele) and that harbours HPV16 integrations (minor allele). Sum of absolute offset of segments and junctions are shown. Unit-cycles (shadowed areas) in LGM are denoted with repeat time. The frame enclosed copy number is corresponding to unit-cycle (the NO.4) only containing the segments in solid frame. The unit-cycles might exist in the form of circular ecDNA.

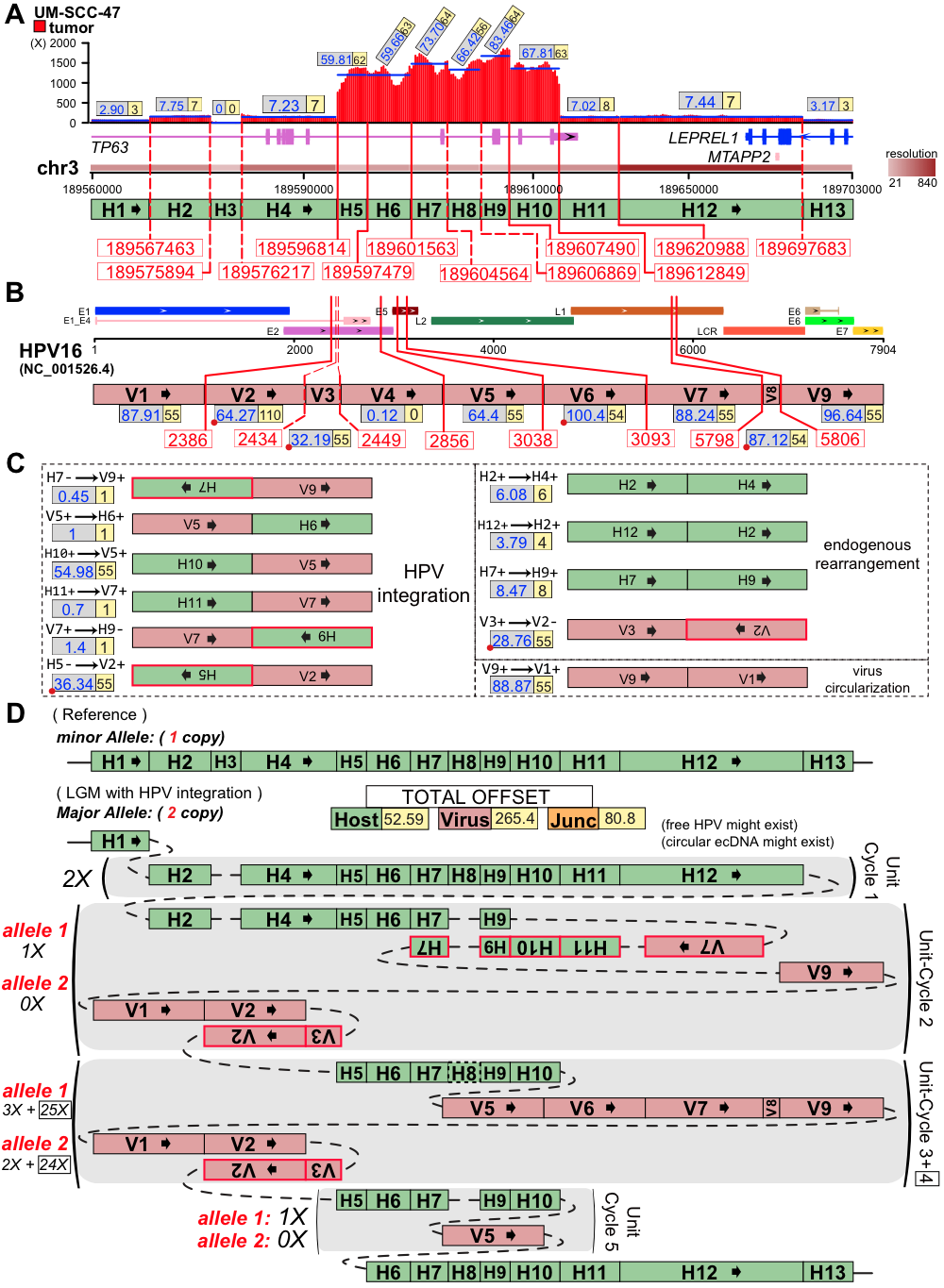

**Supplementary Figure S9. Presentative LGM at HPV integration sites (gene *TP63*) on chr3 (major allele) of the UM-SCC-47 cell line** [2]. (**A**) Human genomic region flanking HPV integrations are divided into 13 segments (H1∼H13) by VITs (red solid-line) and SVs (red dashed-line) denoted with breakpoints. Depth spectrum is displayed with original (grey frame) and ILP-adjusted (yellow frame) copy numbers of segments. (**B**) Segmentation (V1∼V9) of HPV16 genome by VITs and SVs. The segments less than 100bp are marked with red dot. (**C**) Variant segment junctions utilized in Conjugate graph. (**D**) Resolved alleles of the ‘Simplest LGM’ are indicated as string of coloured segments with copy times, including reference allele (minor allele) and that harbours HPV16 integrations (major allele). Sum of absolute offset of segments and junctions are shown. Unit-cycles (shadowed areas) in LGM are denoted with repeat time. The frame enclosed copy number is corresponding to unit-cycle (the NO.4) only containing the segments in solid frame. Note that the HPV-integrated alleles might have different copies of unit-cycles, which might exist as circular ecDNAs.

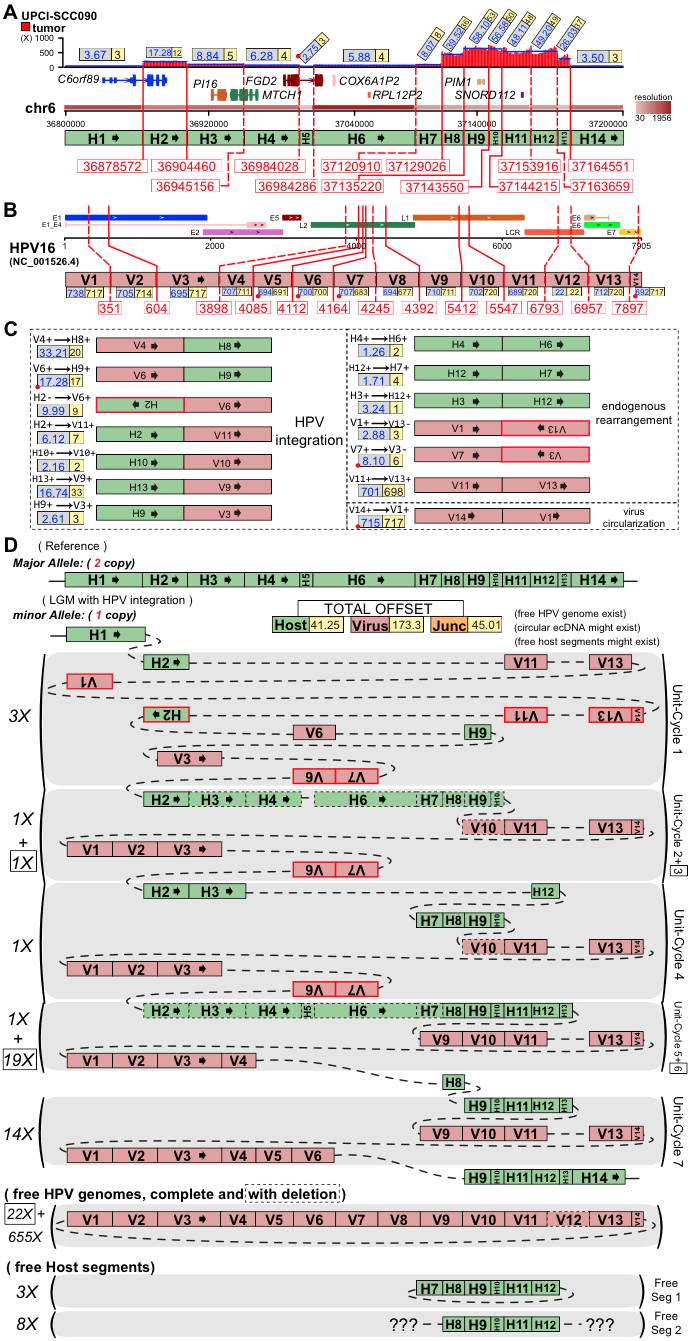

**Supplementary Figure S10.** **Presentative LGM at HPV integration sites (gene *PIM1*) on chr6 (minor allele) on chr6 of the UPCI-SCC090 cell line** [2]. (**A**) Human genomic region flanking HPV integrations are divided into 14 segments (H1∼H14) by VITs (red solid-line) and SVs (red dashed-line) denoted with breakpoints. Depth spectrum is displayed with original (grey frame) and ILP-adjusted (yellow frame) copy numbers of segments. (**B**) Segmentation (V1∼V14) of HPV16 genome by VITs and SVs. The segments less than 100bp are marked with red dot. (**C**) Variant segment junctions utilized in Conjugate graph. (**D**) Resolved alleles of the ‘Simplest LGM’ are indicated as string of coloured segments with copy times, including reference allele (major allele) and that harbours HPV16 integrations (minor allele). Sum of absolute offset of segments and junctions are shown. Unit-cycles (shadowed areas) in LGM are denoted with repeat time. Note that both free HPV genomes and host segments exist with circular and linear structures. The frame enclosed copy number is corresponding to the unit-cycle only containing the segments in solid frame.

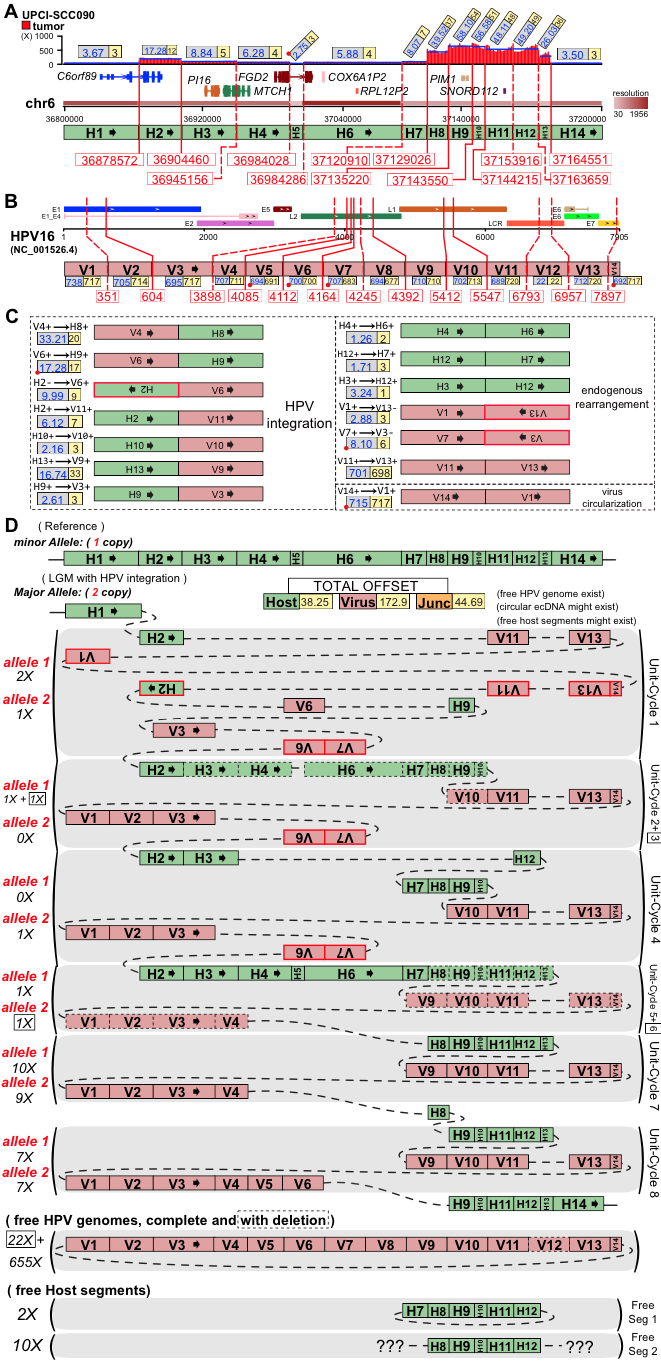

**Supplementary Figure S11.** **Presentative LGM at HPV integration sites (gene *PIM1*) on chr6 (major allele) of the UPCI-SCC090 cell line** [2]. (**A**) Human genomic region flanking HPV integrations are divided into 14 segments (H1∼H14) by VITs (red solid-line) and SVs (red dashed-line) denoted with breakpoints. Depth spectrum is displayed with original (grey frame) and ILP-adjusted (yellow frame) copy numbers of segments. (**B**) Segmentation (V1∼V14) of HPV16 genome by VITs and SVs. The segments less than 100bp are marked with red dot. (**C**) Variant segment junctions utilized in Conjugate graph. (**D**) Resolved alleles of the ‘Simplest LGM’ are indicated as string of coloured segments with copy times, including reference allele (minor allele) and that harbours HPV16 integrations (major allele). Sum of absolute offset of segments and junctions are shown. Unit-cycles (shadowed areas) in LGM are denoted with repeat time. Note that both free HPV genomes and host segments exist with circular and linear structures. The frame enclosed copy number is corresponding to the unit-cycle only containing the segments in solid frame.

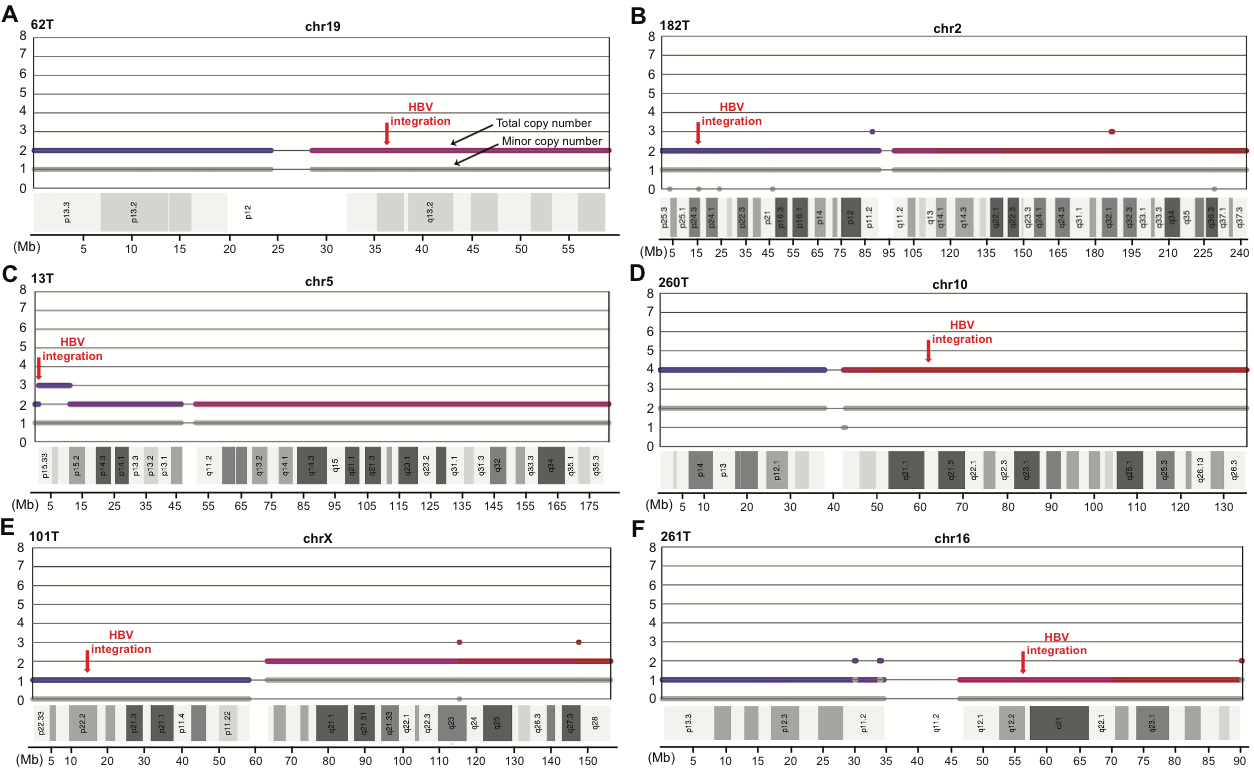

**Supplementary Figure S12.** **Copy number distribution along the HBV-integrated chromosome in six HCC samples** [11]. Figures are from patchwork results. HBV integration sites are denoted by red arrows. Colored and grey bold lines indicate total and minor copy number respectively.

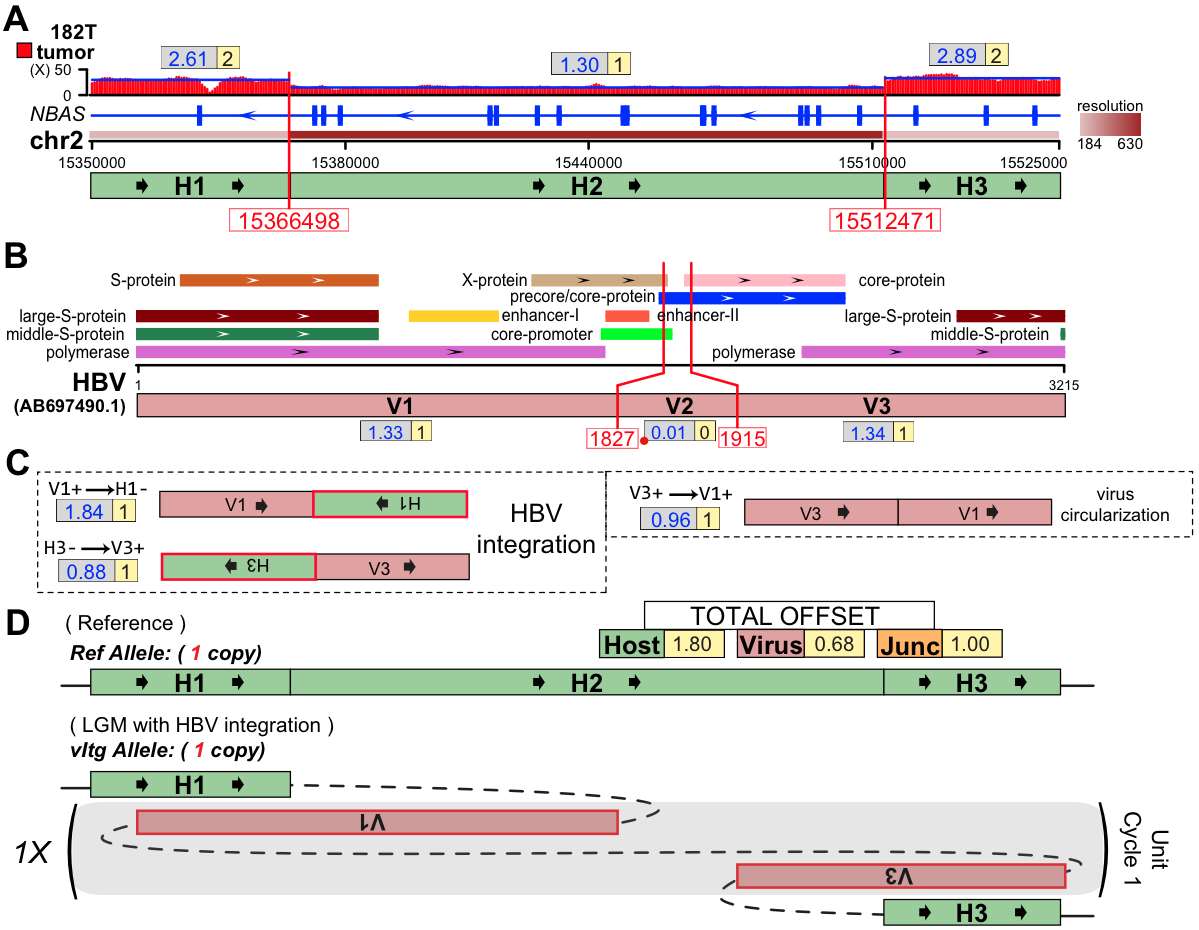

**Supplementary Figure S13.** **Presentative LGM at HBV integration sites (gene *NBAS*) on chr2 of the 182T HCC sample** [11]. (**A**) Human genomic region flanking HBV integrations are divided into three segments (H1∼H3) by VITs denoted with breakpoints. Depth spectrum is displayed with original (grey frame) and ILP-adjusted (yellow frame) copy numbers of segments. (**B**) Segmentation (V1∼V3) of HBV genome by VITs and SVs. The segment less than 100bp is marked with red dot. (**C**) Variant segment junctions utilized in Conjugate graph. (**D**) Resolved alleles of the ‘Simplest LGM’ are indicated as string of coloured segments with copy times, including reference allele and that harbours HBV integrations. Sum of absolute offset of segments and junctions are shown. Unit-cycles (shadowed areas) in LGM are denoted with repeat time.

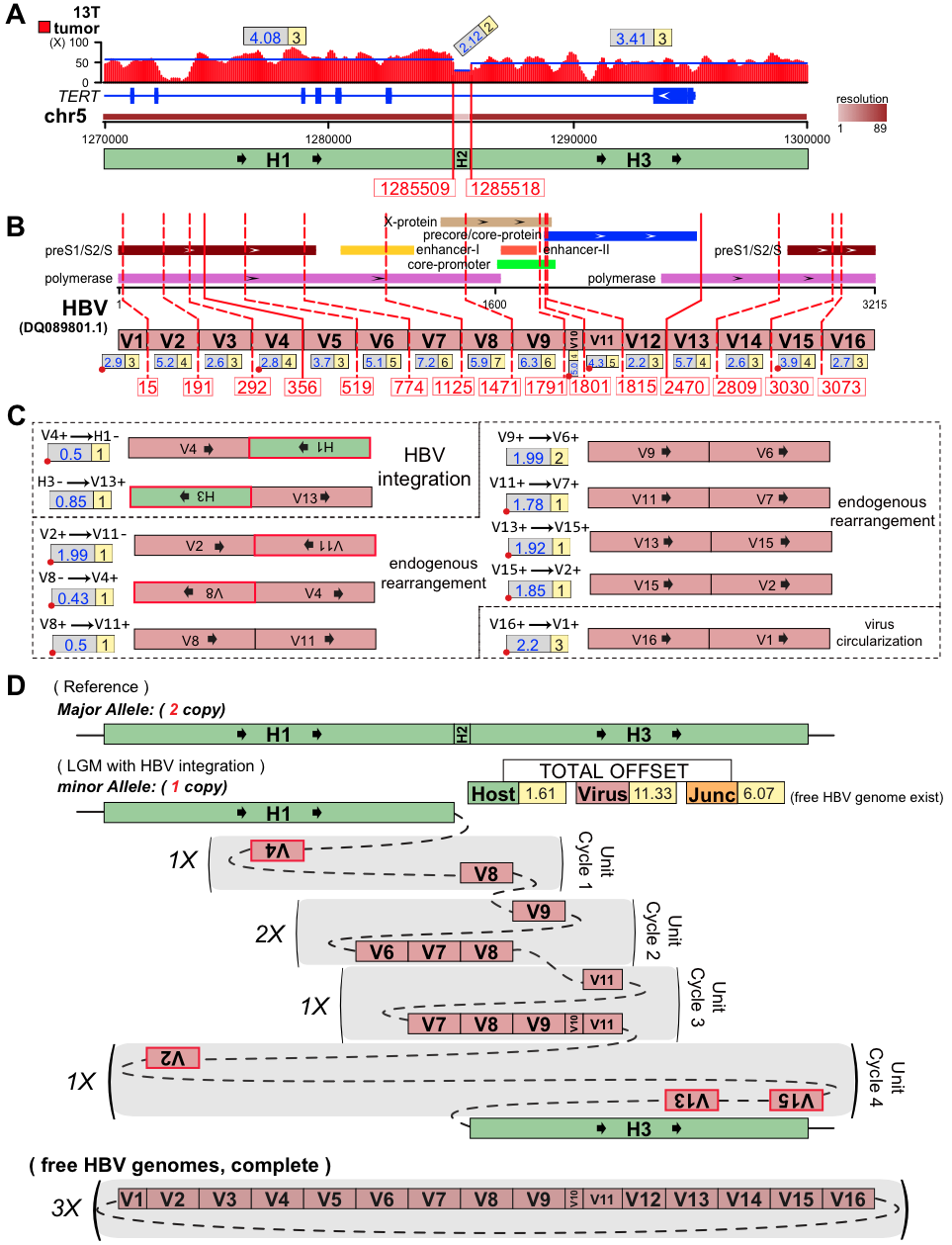

**Supplementary Figure S14.** **Presentative LGM at HBV integration sites (gene *TERT*) on chr5 (minor allele) of the 13T HCC sample** [11]. (**A**) Human genomic region flanking HBV integrations are divided into three segments (H1∼H3) by VITs denoted with breakpoints. Depth spectrum is displayed with original (grey frame) and ILP-adjusted (yellow frame) copy numbers of segments. (**B**) Segmentation (V1∼V16) of HBV genome by VITs and SVs. The segment less than 100bp is marked with red dot. (**C**) Variant segment junctions utilized in Conjugate graph. (**D**) Resolved alleles of the ‘Simplest LGM’ are indicated as string of coloured segments with copy times, including reference allele (major allele) and that harbours HBV integrations (minor allele). Sum of absolute offset of segments and junctions are shown. Unit-cycles (shadowed areas) in LGM are denoted with repeat time. Note that the free HBV genomes might exist.

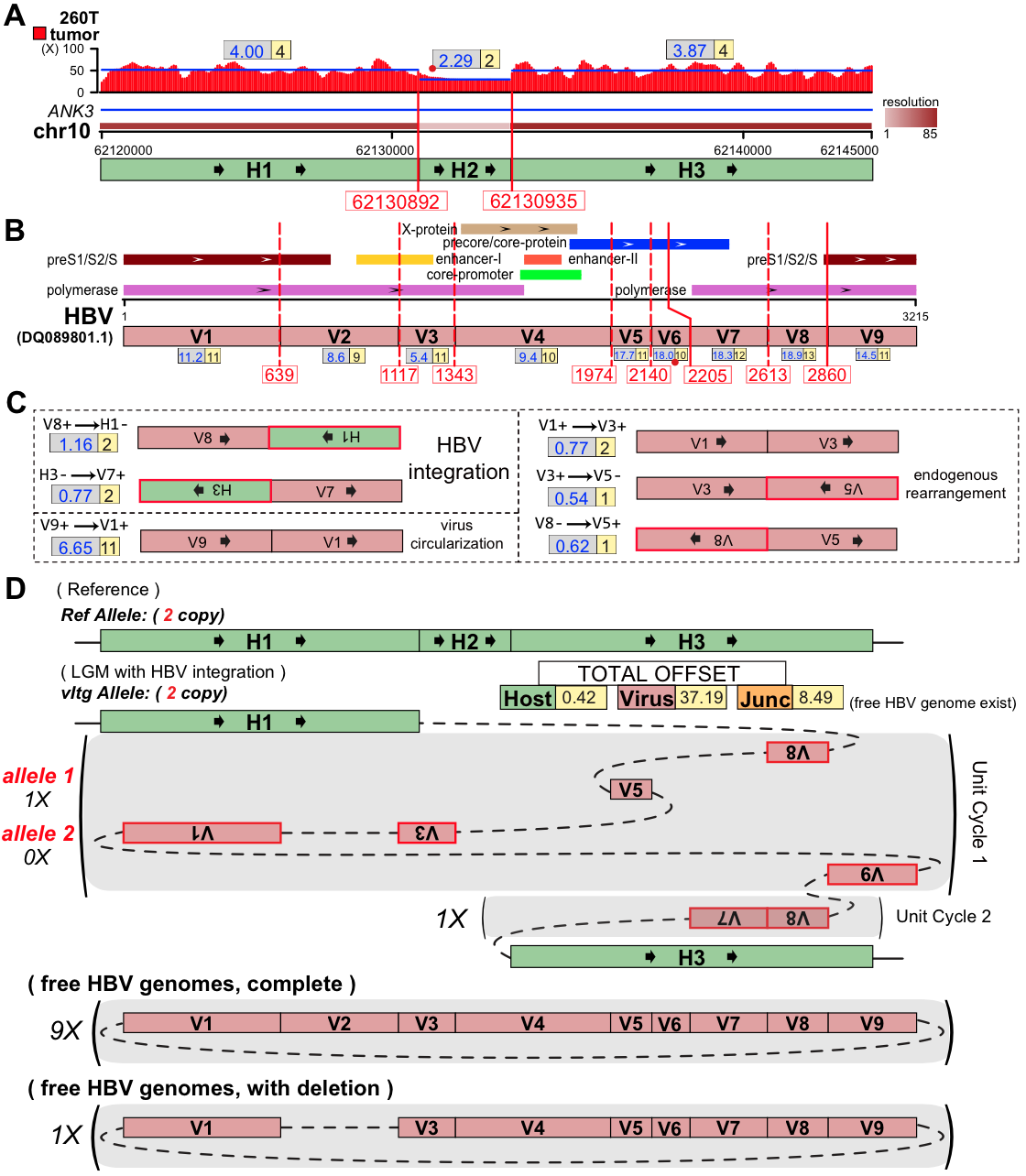

**Supplementary Figure S15.** **Presentative LGM at HBV integration sites (gene *ANK3*) on chr10 of the 260T HCC sample** [11]. (**A**) Human genomic region flanking HBV integrations are divided into three segments (H1∼H3) by VITs denoted with breakpoints. Depth spectrum is displayed with original (grey frame) and ILP-adjusted (yellow frame) copy numbers of segments. (**B**) Segmentation (V1∼V9) of HBV genome by VITs and SVs. The segment less than 100bp is marked with red dot. (**C**) Variant segment junctions utilized in Conjugate graph. (**D**) Resolved alleles of the ‘Simplest LGM’ are indicated as string of coloured segments with copy times, including reference allele and that harbours HBV integrations. Sum of absolute offset of segments and junctions are shown. Unit-cycles (shadowed areas) in LGM are denoted with repeat time. Note that the HBV-integrated alleles might have different copies of unit-cycles, which might exist as circular ecDNA. The HBV free genome (both complete and with deletion) might exist.

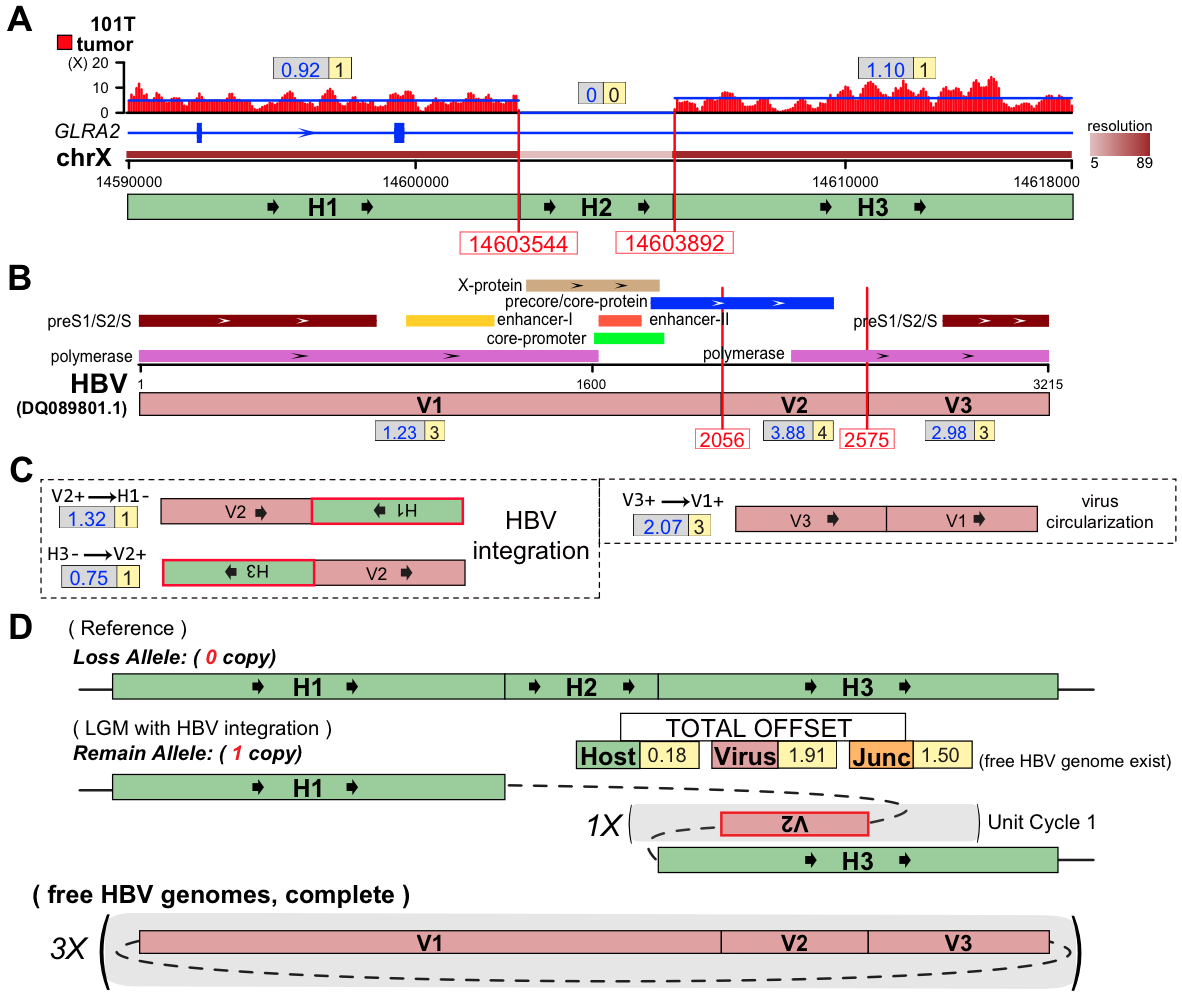

**Supplementary Figure S16.** **Presentative LGM at HBV integration sites (gene *GLRA2*) on chrX of the 101T HCC sample** [11]. (**A**) Human genomic region flanking HBV integrations are divided into three segments (H1∼H3) by VITs denoted with breakpoints. Depth spectrum is displayed with original (grey frame) and ILP-adjusted (yellow frame) copy numbers of segments. (**B**) Segmentation (V1∼V3) of HBV genome by VITs. (**C**) Variant segment junctions utilized in Conjugate graph. (**D**) Resolved alleles of the ‘Simplest LGM’ are indicated as string of coloured segments with copy times, including reference allele (loss allele) and that harbours HBV integrations (remain allele). Sum of absolute offset of segments and junctions are shown. Unit-cycles (shadowed areas) in LGM are denoted with repeat time. Note that the free HBV genomes might exist.

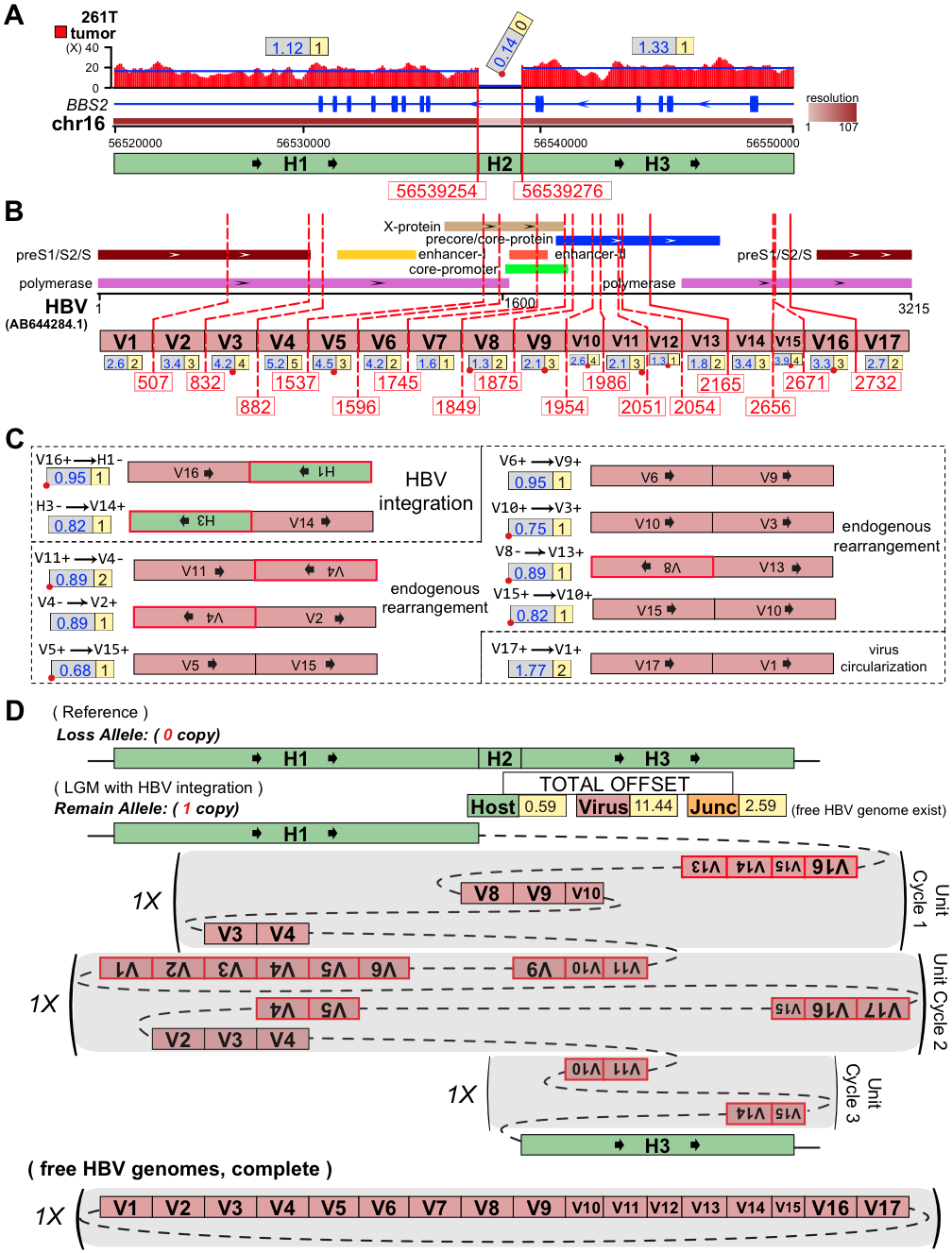

**Supplementary Figure S17.** **Presentative LGM at HBV integration sites (gene *BBS2*) on chr16 of the 261T HCC sample** [11]. (**A**) Human genomic region flanking HBV integrations are divided into three segments (H1∼H3) by VITs denoted with breakpoints. Depth spectrum is displayed with original (grey frame) and ILP-adjusted (yellow frame) copy numbers of segments. (**B**) Segmentation (V1∼V17) of HBV genome by VITs and SVs. The segment less than 100bp is marked with red dot. (**C**) Variant segment junctions utilized in Conjugate graph. (**D**) Resolved alleles of the ‘Simplest LGM’ are indicated as string of coloured segments with copy times, including reference allele (loss allele) and that harbours HBV integrations (remain allele). Sum of absolute offset of segments and junctions are shown. Unit-cycles (shadowed areas) in LGM are denoted with repeat time. Note that the free HBV genomes might exist.

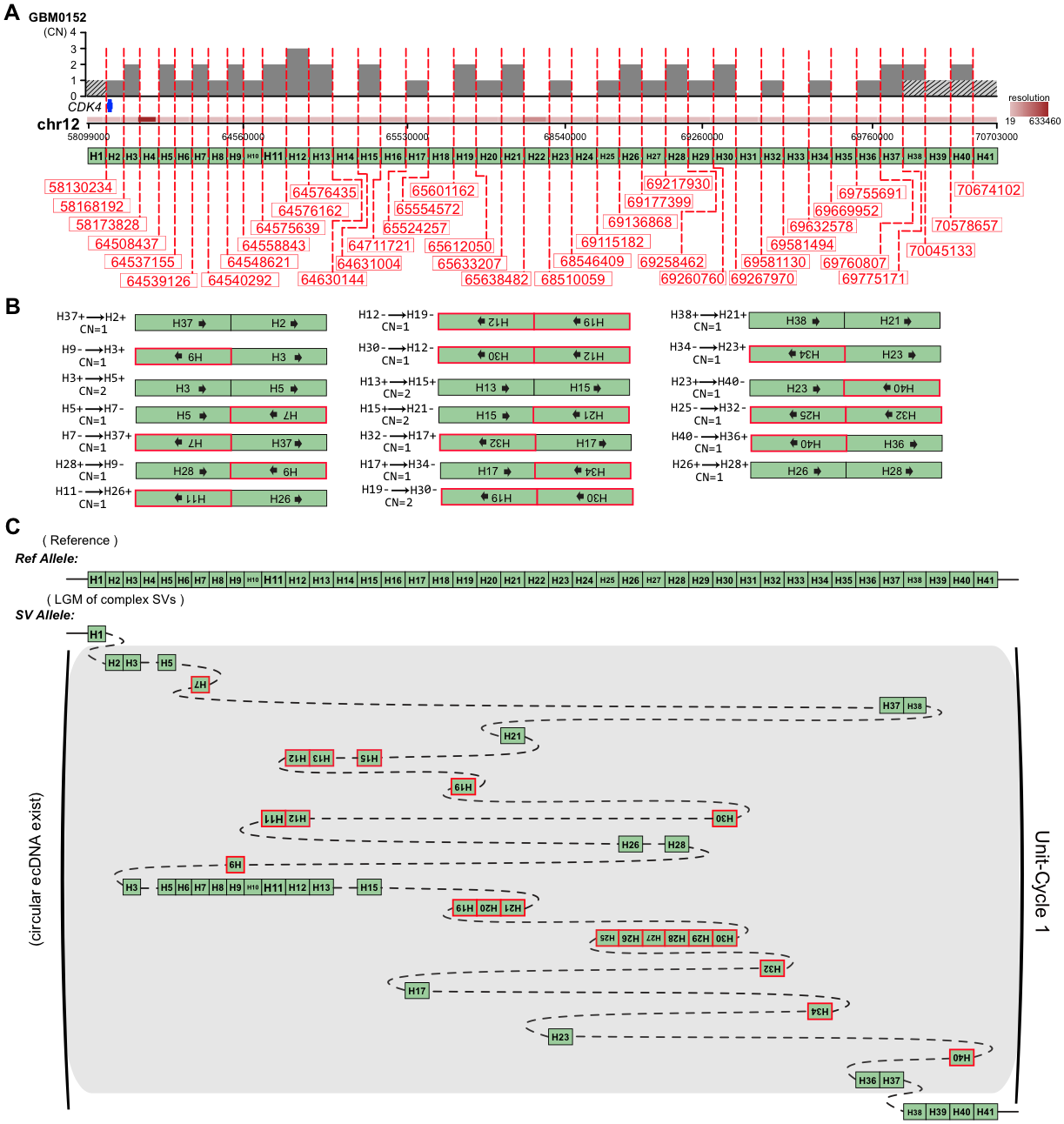

**Supplementary Figure S18.** **Presentative LGM at complex SV locus on chr12 of the GBM0152 cancer sample** [15]. (**A**) Human genomic region is divided into 41 segments (H1∼H41) by SVs denoted with breakpoints. Depth spectrum is displayed as copy numbers of segments. (**B**) Variant segment junctions utilized in Conjugate graph. (**C**) Resolved alleles of the ‘Simplest LGM’ are indicated as string of coloured segments with copy times, including reference allele and SVs LGM allele. Unit-cycles (shadowed areas) in LGM are denoted with repeat time. Note that the unit-cycle NO.1 might exist as circular ecDNA. In this figure, we only use one copy of the unit-cycle according to the copy number shown in the Figure.6D-F in Yang et al. 2013 Cell [15]. The depth specturms with slash shadow are added for *source* (H1) and *sink* (H38,H39,H40,H41) of the LGM in our algorithm.

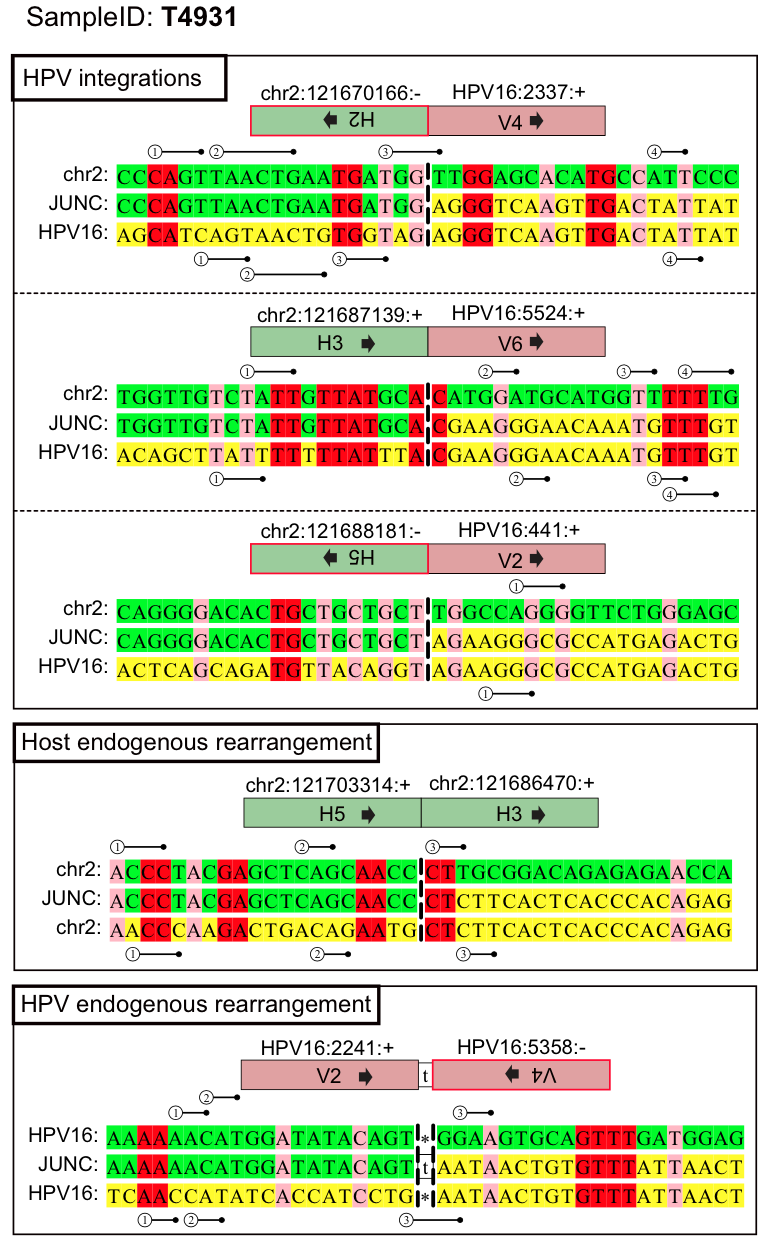

**Supplementary Figure S19.** **Alignment of the sequence around the integration site between the human genome and the HPV16 genome, and the endogenous rearrangements on human genome and the HPV16 genome respectively, in the T4931 sample** [7]. The junction boundaries are shown as vertical dashed lines. All viral sequences are from the reference strand. Green, upstream junction partner; yellow, downstream junction partner; red, nucleotides that vertically align to both reference sequences (aligned microhomologous bases); pairwise numbered sticks, slipped microhomologous bases. All microhomologous bases follow the 5’-to-3’ direction. The junction segment IDs are corresponding to the segments in the resolved local genome maps (**Figure 2** and **Supplementary Figure S3**, **Supplementary Table S3**).

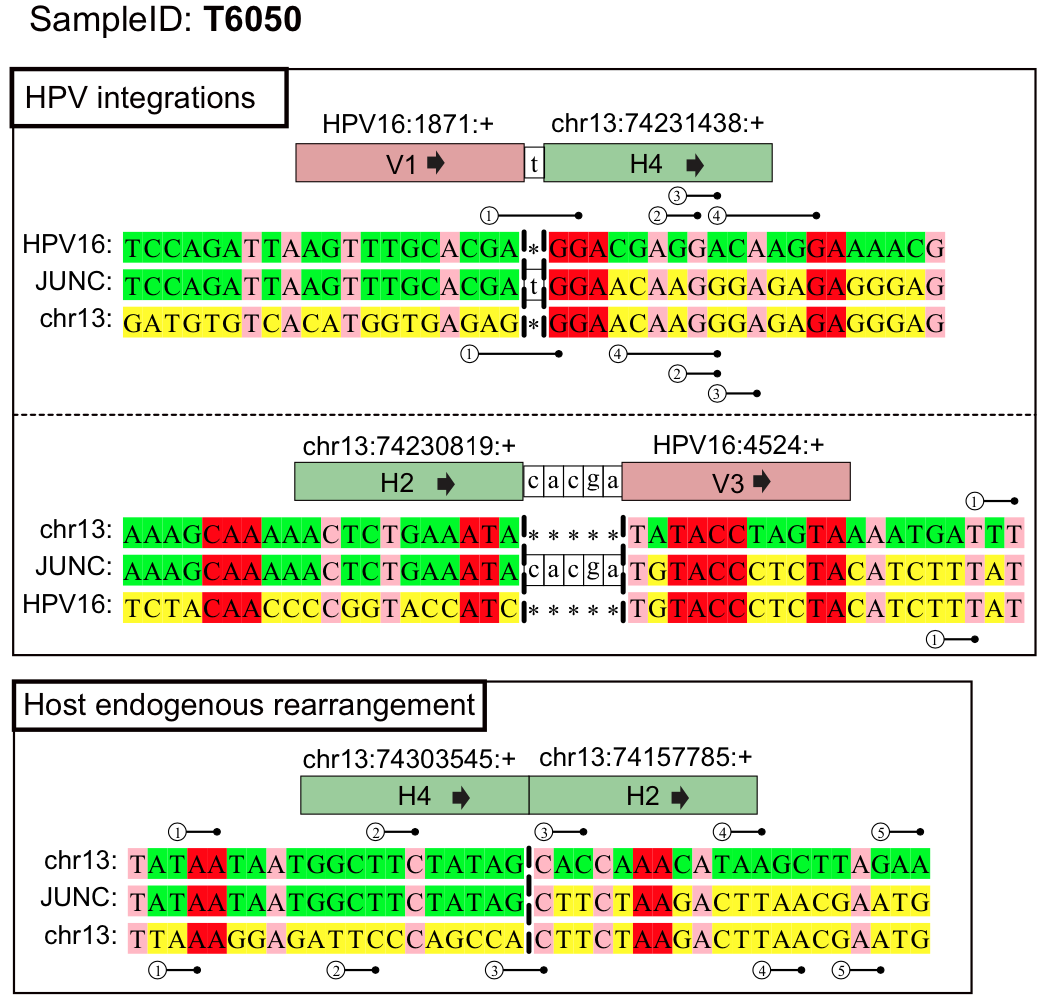

**Supplementary Figure S20.** **Alignment of the sequence around the integration site between the human genome and the HPV16 genome, and the endogenous rearrangements on human genome, in the T6050 sample** [7]. The junction boundaries are shown as vertical dashed lines. All viral sequences are from the reference strand. Green, upstream junction partner; yellow, downstream junction partner; red, nucleotides that vertically align to both reference sequences (aligned microhomologous bases); pairwise numbered sticks, slipped microhomologous bases. All microhomologous bases follow the 5’-to-3’ direction. The junction segment IDs are corresponding to the segments in the resolved local genome map (**Supplementary Figure S5**, **Supplementary Table S3**).

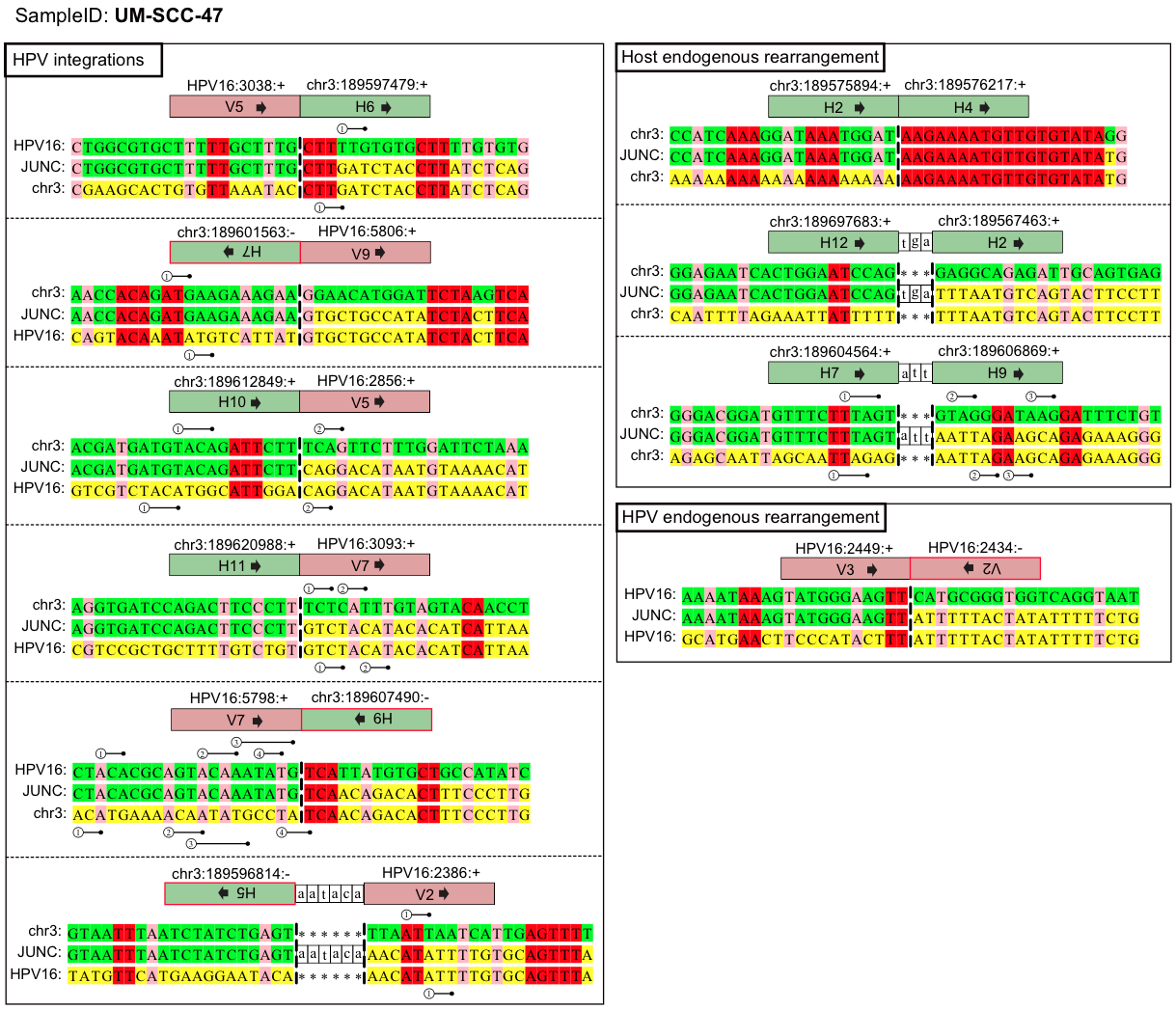

**Supplementary Figure S21.** **Alignment of the sequence around the integration site between the human genome and the HPV16 genome, and the endogenous rearrangements on human genome and the HPV16 genome respectively, in the UM-SCC-47 cell line** [2]. The junction boundaries are shown as vertical dashed lines. All viral sequences are from the reference strand. Green, upstream junction partner; yellow, downstream junction partner; red, nucleotides that vertically align to both reference sequences (aligned microhomologous bases); pairwise numbered sticks, slipped microhomologous bases. All microhomologous bases follow the 5’-to-3’ direction. The junction segment IDs are corresponding to the segments in the resolved local genome maps (**Supplementary Figure S8** and **S9**, **Supplementary Table S4**).

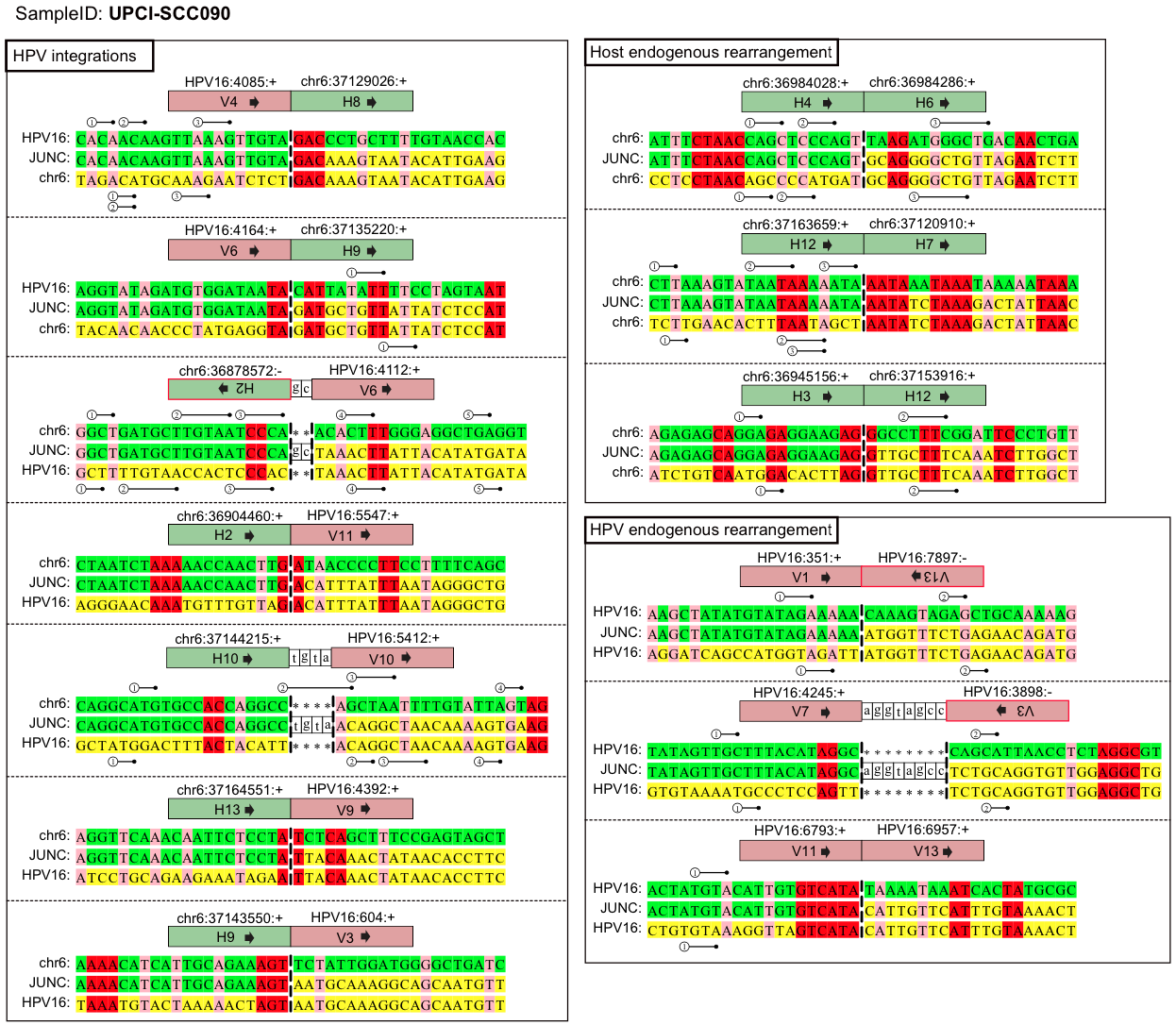

**Supplementary Figure S22.** **Alignment of the sequence around the integration site between the human genome and the HPV16 genome, and the endogenous rearrangements on human genome and the HPV16 genome respectively, in the UPCI-SCC090 cell line** [2]. The junction boundaries are shown as vertical dashed lines. All viral sequences are from the reference strand. Green, upstream junction partner; yellow, downstream junction partner; red, nucleotides that vertically align to both reference sequences (aligned microhomologous bases); pairwise numbered sticks, slipped microhomologous bases. All microhomologous bases follow the 5’-to-3’ direction. The junction segment IDs are corresponding to the segments in the resolved local genome maps (**Supplementary Figure S10** and **S11**, **Supplementary Table S4**).

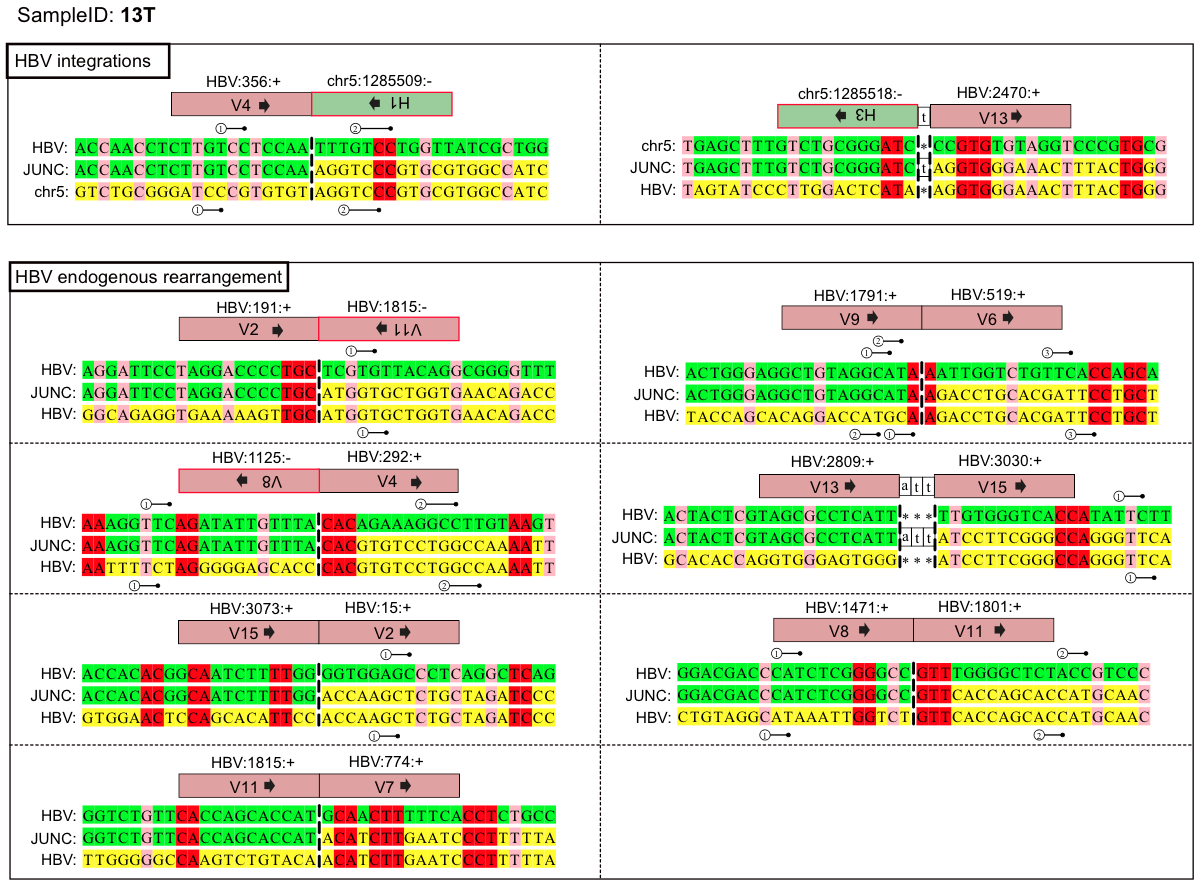

**Supplementary Figure S23.** A**lignment of the sequence around the integration site between the human genome and the HBV genome, and the endogenous rearrangements on HBV genome, in the 13T HCC sample** [11]. The junction boundaries are shown as vertical dashed lines. All viral sequences are from the reference strand. Green, upstream junction partner; yellow, downstream junction partner; red, nucleotides that vertically align to both reference sequences (aligned microhomologous bases); pairwise numbered sticks, slipped microhomologous bases. All microhomologous bases follow the 5’-to-3’ direction. The junction segment IDs are corresponding to the segments in the resolved local genome maps (**Supplementary Figure S14**, **Supplementary Table S6**).

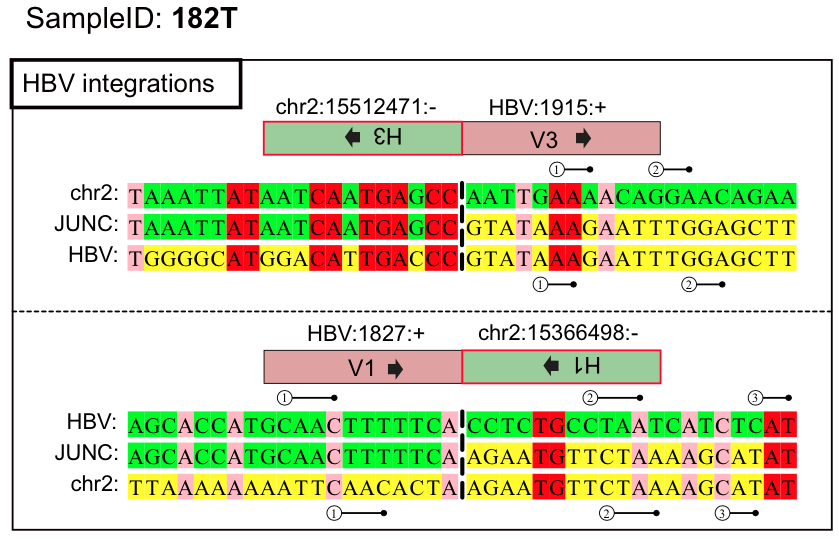

**Supplementary Figure S24.** **Alignment of the sequence around the integration site between the human genome and the HBV genome in the 182T HCC sample** [11]. The junction boundaries are shown as vertical dashed lines. All viral sequences are from the reference strand. Green, upstream junction partner; yellow, downstream junction partner; red, nucleotides that vertically align to both reference sequences (aligned microhomologous bases); pairwise numbered sticks, slipped microhomologous bases. All microhomologous bases follow the 5’-to-3’ direction. The junction segment IDs are corresponding to the segments in the resolved local genome maps (**Supplementary Figure S13**, **Supplementary Table S6**).

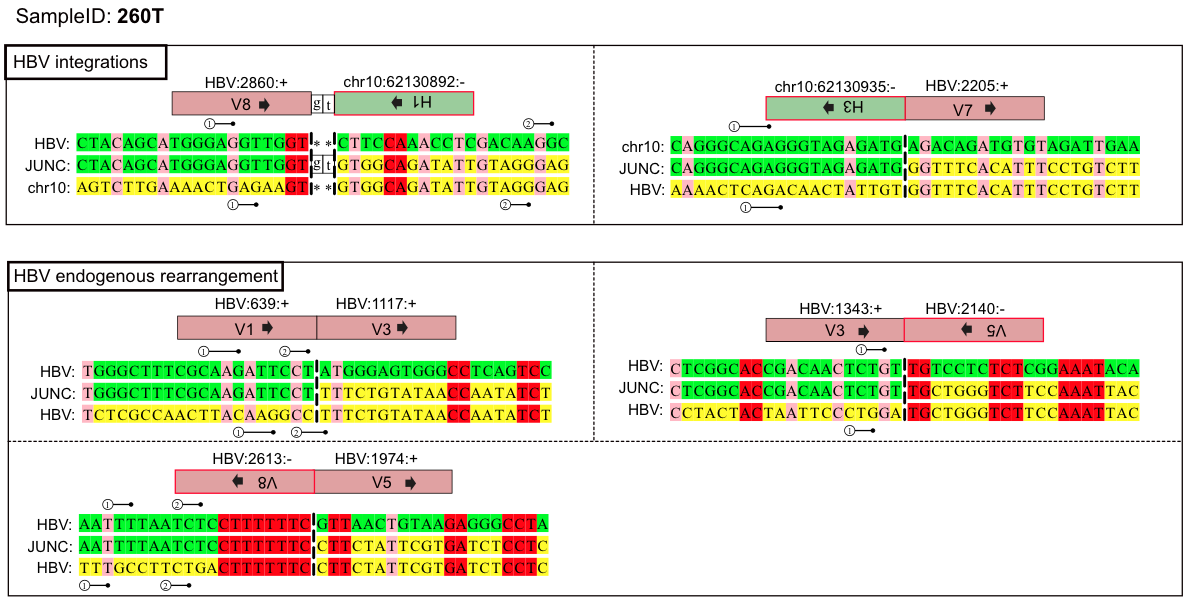

**Supplementary Figure S25.** **Alignment of the sequence around the integration site between the human genome and the HBV genome, and the endogenous rearrangements on HBV genome, in the 260T HCC sample** [11]. The junction boundaries are shown as vertical dashed lines. All viral sequences are from the reference strand. Green, upstream junction partner; yellow, downstream junction partner; red, nucleotides that vertically align to both reference sequences (aligned microhomologous bases); pairwise numbered sticks, slipped microhomologous bases. All microhomologous bases follow the 5’-to-3’ direction. The junction segment IDs are corresponding to the segments in the resolved local genome maps (**Supplementary Figure S15**, **Supplementary Table S6**).

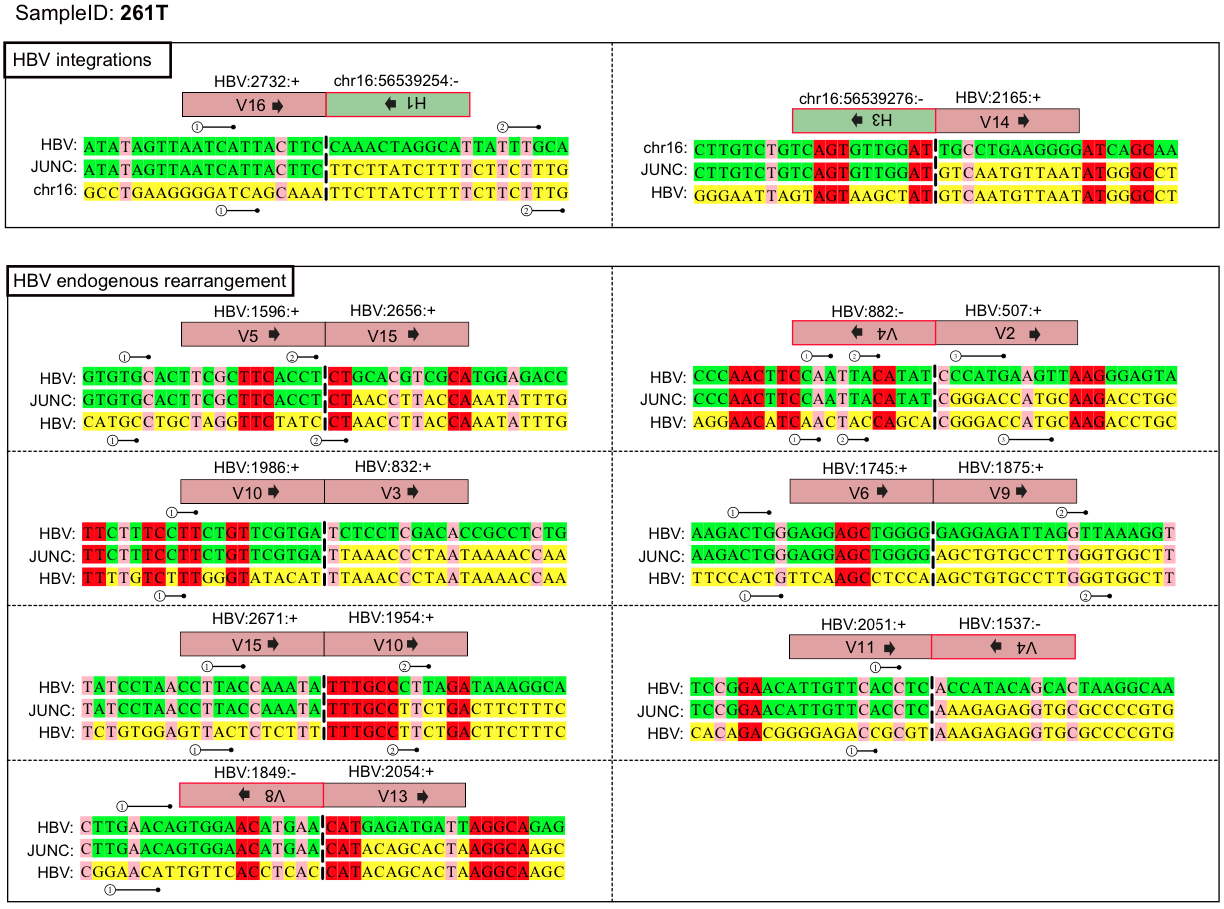

**Supplementary Figure S26.** **Alignment of the sequence around the integration site between the human genome and the HBV genome, and the endogenous rearrangements on HBV genome, in the 261T HCC sample** [11]. The junction boundaries are shown as vertical dashed lines. All viral sequences are from the reference strand. Green, upstream junction partner; yellow, downstream junction partner; red, nucleotides that vertically align to both reference sequences (aligned microhomologous bases); pairwise numbered sticks, slipped microhomologous bases. All microhomologous bases follow the 5’-to-3’ direction. The junction segment IDs are corresponding to the segments in the resolved local genome maps (**Supplementary Figure S17**, **Supplementary Table S6**).

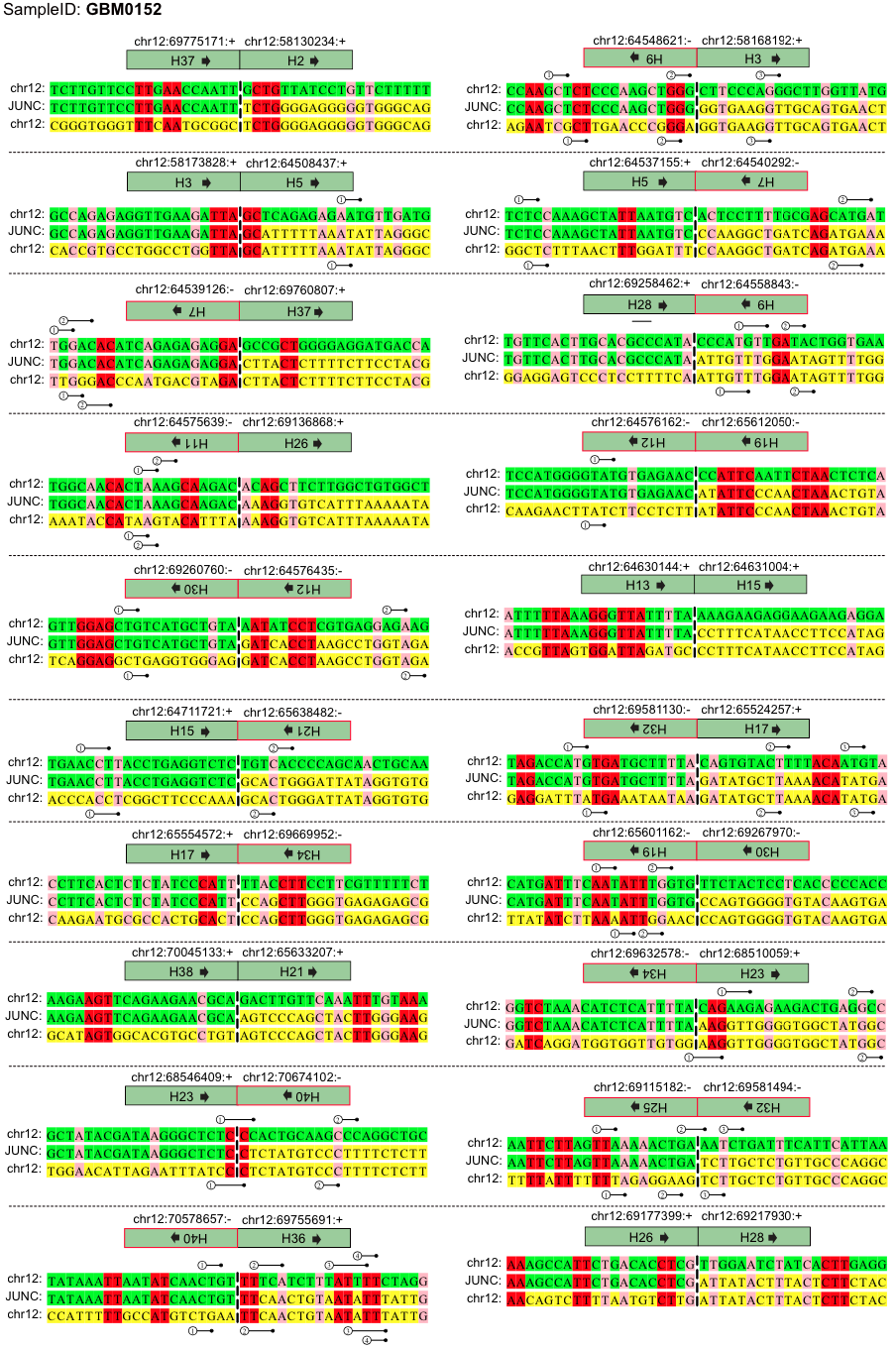

**Supplementary Figure S27.** **Alignment of the sequence around the endogenous complex rearrangements on human genome in the GBM0152 cancer sample** [15]. The junction boundaries are shown as vertical dashed lines. All viral sequences are from the reference strand. Green, upstream junction partner; yellow, downstream junction partner; red, nucleotides that vertically align to both reference sequences (aligned microhomologous bases); pairwise numbered sticks, slipped microhomologous bases. All microhomologous bases follow the 5’-to-3’ direction. The junction segment IDs are corresponding to the segments in the resolved local genome maps (**Supplementary Figure S18**, **Supplementary Table S7**).

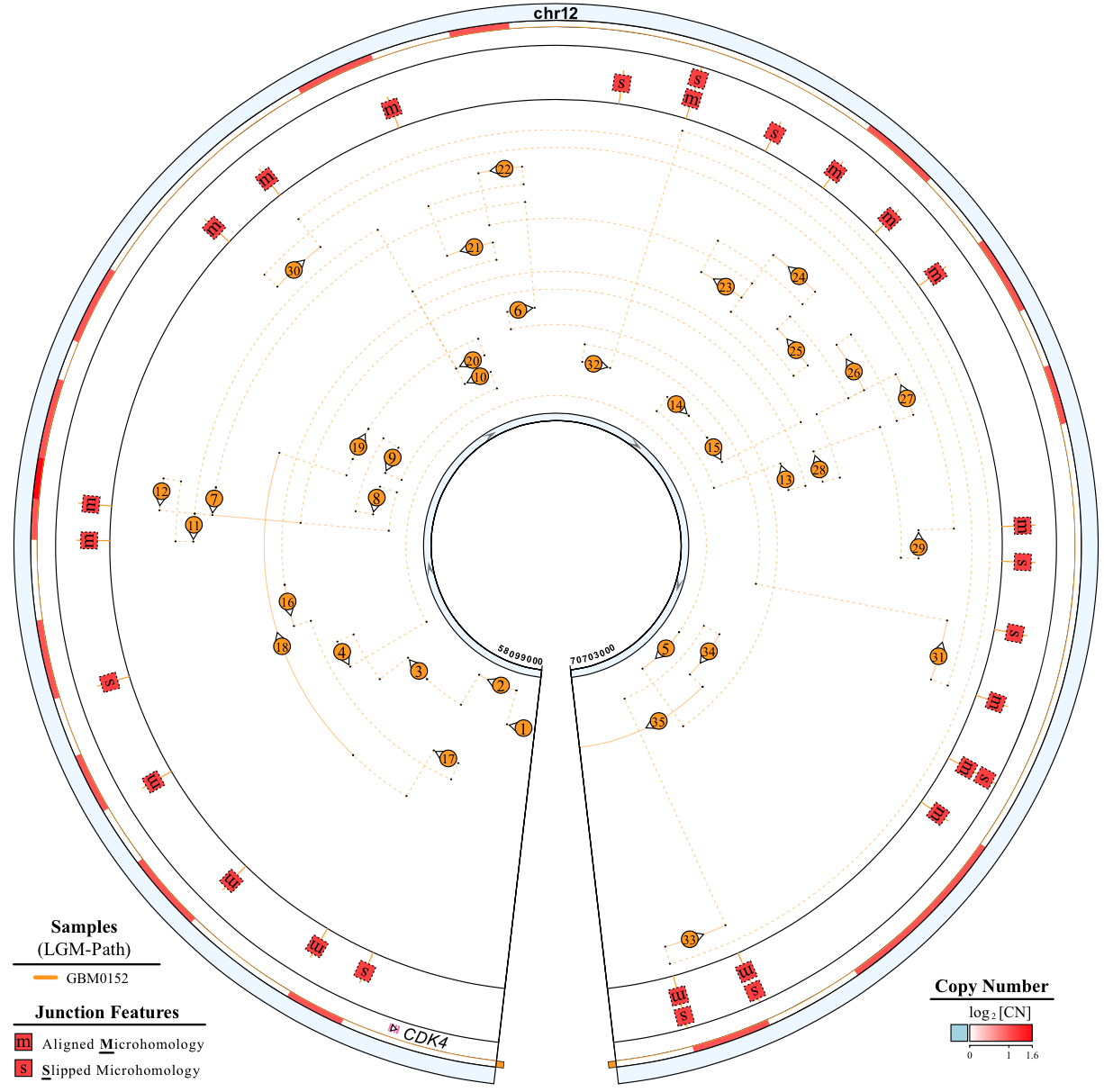

**Supplementary Figure S28.** **Features of the complex SV LGM in GBM0152 sample. Human genomic segments related to complex SVs LGMs are shown as sectors with their relevant LGM path.** Segments of LGMs are denoted by numbered concentric arcs. The numbers, starting from 1, indicate the order in which each arc is visited starting from the source segment of an LGM. Sequential segments in LGM could be merged in single arc (**Supplementary Table S9**). Features of rearrangement sites are depicted as single-letter icons. DNA copy number (CN) is displayed in gradient red colour.

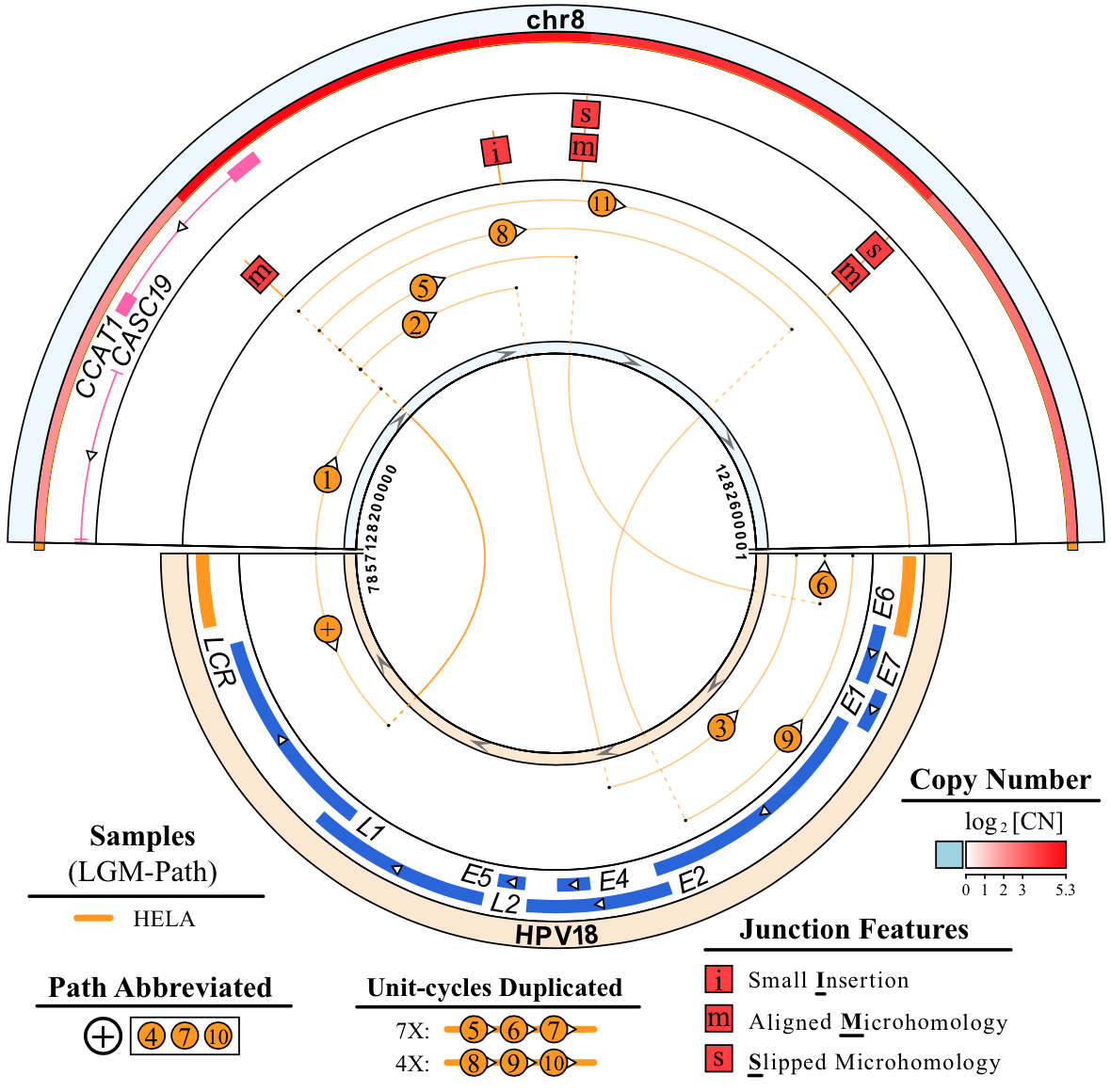

**Supplementary Figure S29.** **Features of HPV18 integrated LGM in HeLa cell line. Human genomic segments are shown as sectors with the LGM path. Segments of LGMs are denoted by numbered concentric arcs.** The numbers, starting from 1, indicate the order in which each arc is visited starting from the source segment of an LGM. Each special symbol shown in “Path Abbreviated” section in the figure legends denotes an arc that is visited more than once in the LGM, such as the ‘+’ symbol for NO.4, NO.7, and NO.10 arcs in the LGM. Sequential segments in LGM could be merged in single arc (**Supplementary Table S9**). Repeat times of unit-cycles in LGMs are stated in figure legend. Features of HPV18 integration sites are depicted as single-letter icons. DNA copy number (CN) is displayed in gradient red colour. The HPV18 genome reference is NC 001357.1 from the NCBI Nucleotide database.
